# A sensitive and accurate framework for population-scale structural variant discovery and genotyping across sequence types

**DOI:** 10.64898/2026.01.10.698766

**Authors:** Guangbao Luo, Li Xiao, Zhangjun Fei, Xin Wang

## Abstract

Structural variation underlies much of the phenotypic and evolutionary diversity. However, accurate discovery and genotyping of structural variants (SVs) at population scale remain challenging due to the varied characteristics of sequencing technologies and the complexity of genome architectures. Here, we introduce PSVGT, a unified framework that integrates short-read and long-read data, de novo assembled contigs, and chromosome-level assemblies to enable comprehensive SV detection and genotyping across diploid, polyploid, and allopolyploid genomes. PSVGT employs an integrated signaling module to extract precise insertion and deletion breakpoints, coupled with the ploidy-aware KLOOK clustering algorithm and local depth-adaptive filtering to resolve multi-allelic events and accommodate the uneven coverage characteristic of complex genomic regions. Benchmarking on simulated and real datasets demonstrates that PSVGT consistently outperforms state-of-the-art tools across sequence types, with advantages particularly in complex genomes and low-coverage long-read data. PSVGT fills a critical gap in scalable SV analysis by leveraging underutilized short-read data and enabling robust characterization of SVs across diverse genome architectures from diploids to polyploids–thereby facilitating population-scale analyses and pan-genome research.

## Background

Genomic variation underlies phenotypic diversity and plays a crucial role in shaping evolution and gene function. Such variation spans a wide spectrum, from single-nucleotide changes to highly complex genomic rearrangements. Large structural variants (SVs), including deletions, insertions, inversions, duplications, and translocations, have profound effects on genomic architecture, altering gene function and expression^1, 2^. Consequently, population-scale studies of SVs have attracted increasing attention. Nonetheless, reliable detection and genotyping of SVs at population scale remain challenging. Advances in high-throughput sequencing have enabled the sequencing of thousands of individuals, leading to landmark initiatives such as the Human 1000 Genomes Project, the Arabidopsis 1001 Genomes Project, and the 3000 Rice Genomes Project. These efforts have produced extensive catalogs of genetic variation, greatly advancing pan-genomics, genome-wide association studies (GWAS), population genetics, and domestication research. However, most analyses have focused predominantly on single nucleotide polymorphisms (SNPs), which primarily rely on aligning short reads to a linear reference genome.

Over the past two decades, next-generation sequencing (NGS) has greatly facilitated the study of SVs. Consequently, a wide range of computational tools for SV identification have been developed. Nonetheless, these short-read-based methods often suffer from high false-positive and false-negative rates^5, 6^. Tools such as DELLY^7^, LUMPY^8^, and Manta^8^ share similar principles by integrating paired-end, split-read, and/or read-depth signals. However, the intrinsic limitations of short reads‒and the even shorter fragments derived from split reads‒often lead to alignment errors and unreliable SV calls. The emergence of third-generation long-read sequencing technologies has substantially improved SV detection. Recent tools such as Sniffles2^9^, cuteSV^10^, VolcanoSV^11^, and DeBreak^12^ leverage long reads to resolve complex SVs with higher accuracy. However, the high cost of long-read sequencing remains a major barrier to population-scale studies, while vast amounts of existing short-read data remain underutilized.

At the same time, advances in sequencing and bioinformatics have enabled the generation of increasingly complete genome assemblies, some achieving telomere-to-telomere (T2T) completeness. Because assembled contigs are substantially longer and more accurate than raw reads, they enable the identification of complex rearrangements. Several tools, including AsmVar^14^, Assemblytics^15^, and SyRI^16^, have been developed to leverage genome assemblies for SV discovery. For haplotype-resolved or phased genome assemblies, dipcall^17^ and SVIM-asm^18^ enable accurate SV detection in diploid species. However, SV detection and genotyping in haplotype-resolved assemblies of polyploid species‒such as rose^19^, potato^20^, and sweetpotato^21^‒have not yet been reported. More recently, graph-based pangenomes have emerged as a powerful framework for representing genome-wide variation within a species, overcoming limitations of a single linear reference. Graph-based approaches can encode multiple haplotypes and SVs simultaneously, and effective graph-based pangenome construction depends on integrating diverse data types, including high-quality, haplotype-resolved assemblies from long-read sequencing to capture complex SVs, together with short-read datasets to refine variant breakpoints and genotypes. Despite these advances, current SV detection methods remain largely sequencing-platform specific, and no existing tool supports comprehensive SV identification and genotyping across short reads, long reads, and haplotype-resolved assemblies.

To address these systemic challenges, we developed PSVGT, an integrated framework for population-scale SV detection and genotyping across short reads, long reads, assembled contigs, and chromosome-level assemblies. PSVGT incorporates a novel KLOOK algorithm, which leverages window-based signals and local depth adaptation to efficiently resolve multiple alleles within the same genomic region. Benchmark analyses demonstrate that PSVGT outperforms existing tools in both recall and genotyping accuracy, especially for low-depth long reads and haplotype-resolved assemblies, while also delivering high-precision SV calls from short-read datasets. In addition, the genotyping modules in PSVGT support multiple sequence types, enabling rapid targeted SV genotyping at scale without reprocessing genome-wide SV signals. PSVGT also provides annotations for insertion and deletion (InDel) SVs and includes an automated primer-design module that selects optimal InDel markers for agarose gel electrophoresis analyses, thereby accelerating genetic mapping, marker-assisted selection, and population genetic studies. Collectively, PSVGT addresses the evolving demands of next-generation and long-read genomics, as well as the increasing availability of haplotype-resolved assemblies, by combining comprehensive variant detection and genotyping with marker development in a unified, high-throughput framework.

## Results

### Overview of PSVGT

PSVGT is designed to accommodate a wide range of sequencing data inputs, including Illumina short reads and long reads from PacBio HiFi, PacBio CLR, and ONT platforms in FASTQ format, and *de novo* assembled contigs and chromosome-level genomes in FASTA format, or pre-aligned BAM files from any of these sequence types. Short reads are first assembled into contigs, after which all input sequences are used for SV signal extraction and genotyping. PSVGT provides two modes of SV signal detection: (i) capturing large InDels signals from intra alignments, and (ii) detecting InDels from both inter- and intra-alignments, as well as duplications, inversions, and translocations from inter-alignments (**Fig. 1a**). Candidate SV signals are subsequently processed using the KLOOK clustering algorithm to resolve multi-allelic events, together with local coverage-adaptive filtering to ensure both precision and computational efficiency, particularly in complex genomic regions with uneven sequencing coverage (**Fig. 1b**).

**Fig. 1.**
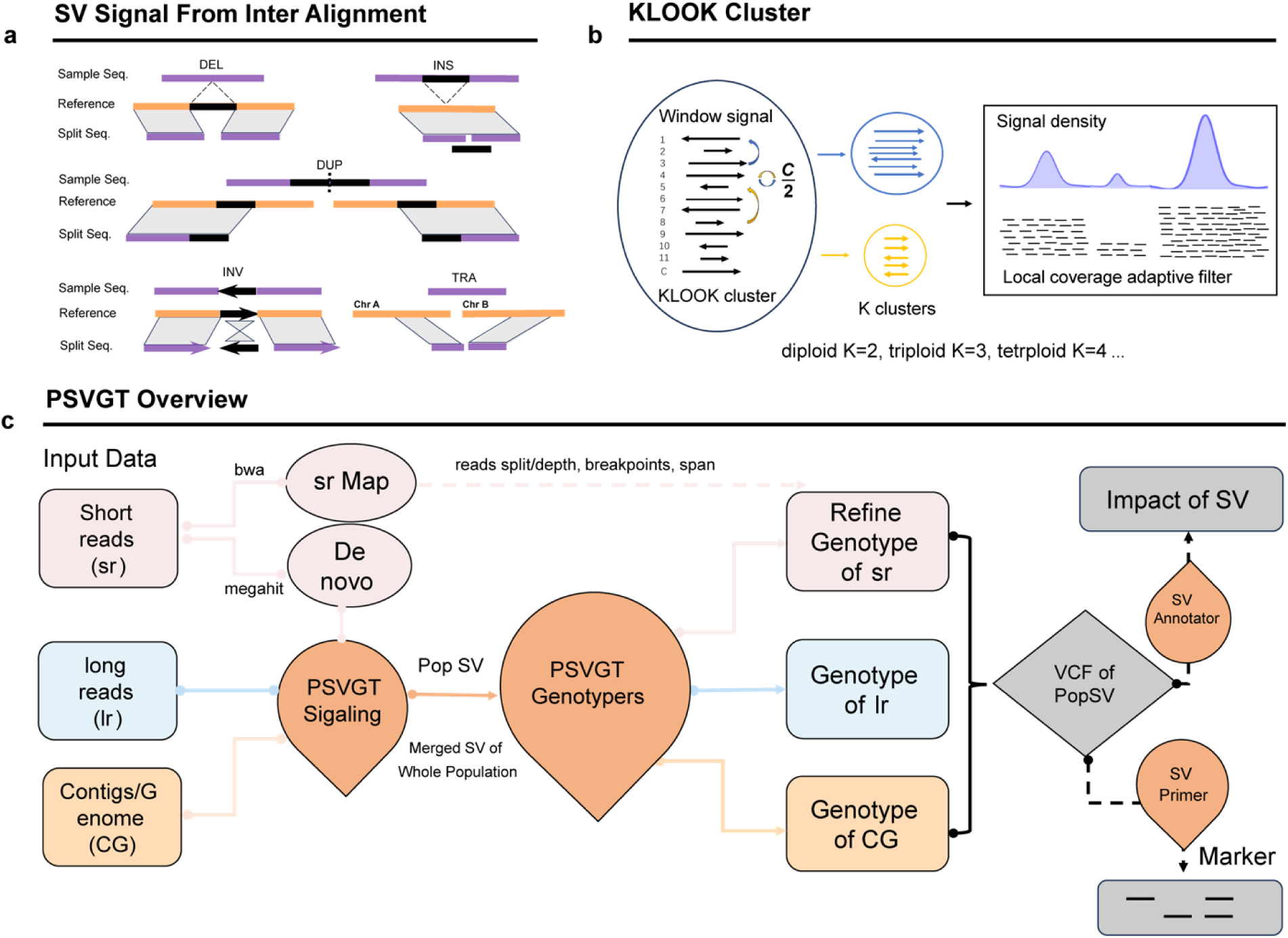
Overview of the PSVGT workflow. **(a)** SV inference from inter-alignments. SVs, including deletions (DEL), insertions (INS), duplications (DUP), inversions (INV), and translocations (TRA), are detected by aligning sample sequences (purple) to the reference genome (orange). Detected SVs are marked in black. (**b)** KLOOK clustering algorithm implemented in PSVGT. Signals within a window are clustered into K groups, where K is defined by ploidy, based on SV start positions, sizes, and local read depth. (**c)** Overall PSVGT workflow. PSVGT accepts diverse types of DNA sequences, including short reads (sr), long reads (lr), and assembled contigs/genomes (CG). Short reads are aligned to the reference genome using BWA and simultaneously de novo assembled into contigs. Candidate SV signals are collected from all samples and processed through KLOOK clustering and local depth-adaptative filtering within the PSVGT Signaling module. Population-level SVs (PopSVs) are then genotyped in each sample using sequence-type-specific sub-modules. Genotype calls from short-read assemblies are further refined using breakpoint information from read alignments. The final population-scale SVs in VCF format support downstream analyses, including marker development via the SV primer module and functional impact annotation via the SV annotation module.

The PSVGT workflow comprises the following steps:

Step 1. *De novo* assembly of short-read datasets using MEGAHIT^22^, followed by mapping all reads and contigs from all samples to the reference genome using minimap2^23^, unless pre-aligned BAM files are provided.

Step 2. Extraction of SV signatures–including CIGAR-based InDels, split reads, spanning reads, breakpoints–from BAM alignment files, followed by KLOOK clustering to resolve multiple-allelic SVs and local depth-based filtering to group and refine candidate SV signals.

Step 3. Merging of candidate SVs across all samples to generate a non-redundant SV set (for sample sizes ≥ 2).

Step 4. Genotyping of candidate SVs in each sample using integrated evidence from CIGAR-based InDel signatures, breakpoints, spanning reads, and split reads.

Step 5. Optionally, refinement of genotypes for *de novo*-assembled or short-read samples using short-read mapping evidence to reduce false positives introduced by assembly errors.

Step 6. Annotation of SVs with genomic coordinates, neighboring gene context, and predicted functional impacts.

Optional: Development of PCR-based SV and CAPS markers for agarose gel electrophoresis or marker-association studies.

A schematic overview of the PSVGT pipeline is shown in **Fig. 1c**, with details provided in the “Materials and Methods” section.

### PSVGT performance on simulated long-read datasets

To evaluate PSVGT performance on long-read data, we simulated multiple human sequencing data types, including haplotype-resolved assemblies, PacBio CLR, PacBio HiFi, and ONT reads. The simulated SV set comprised 10,000 deletions, 10,000 insertions, 1,000 inversions, and 1,000 duplications, following methods adapted from the DeBreak study^12^, along with 365 translocations based on cuteSV benchmarks^10^ (**Fig. S1 & Table S2**). SV size distributions closely matched those observed in the human HG002 genome. The simulated SVs were embedded into the human haplotype-resolved reference genome (hs37d5.fa). Two-thirds of the simulated deletions, insertions, duplications, and inversions were modeled as heterozygous. For translocations, two independent simulated datasets were generated, one containing exclusively homozygous translocations and the other exclusively heterozygous translocations. We benchmarked PSVGT against three state-of-the-art long-read SV callers–DeBreak, cuteSV, and Sniffles2–across all sequencing data types. PSVGT achieved the highest or comparable F1 scores across all SV classes (**Table 1**). Specifically, PSVGT reached overall F1 scores of 99.72% (HiFi), 99.45% (CLR), and 99.67% (ONT), compared with 98.81%, 98.64%, and 98.66% for Sniffles2; 98.78%, 98.30%, and 96.26% for cuteSV, and 99.53%, 99.25%, and 99.36% for DeBreak. PSVGT also consistently delivered high recall and precision across all SV classes (**Table 1**). Notably, PSVGT showed particularly strong performance in resolving complex SVs, achieving recall rates of 99.9% (HiFi), 99.7% (ONT), and 99.6% (CLR) for duplications, substantially exceeding those of Sniffles2 (87.0-89.2%), cuteSV (69.4-75.0%), and DeBreak (95.3-97.8%).

**Table 1.** SV discovery and genotyping accuracy (%) on simulated long-read sequencing data.

| Seq Type | PSVGT |  |  |  | Sniffles2 |  |  |  | cuteSV |  |  |  | DeBreak |  |  |  |
| --- | --- | --- | --- | --- | --- | --- | --- | --- | --- | --- | --- | --- | --- | --- | --- | --- |
|  | Rec | Pre | F1 | GT | Rec | Pre | F1 | GT | Rec | Pre | F1 | GT | Rec | Pre | F1 | GT |

| HiFi |  |  |  |  |  |  |  |  |  |  |  |  |  |  |  |  |
| --- | --- | --- | --- | --- | --- | --- | --- | --- | --- | --- | --- | --- | --- | --- | --- | --- |
| DEL | **99.97** | 99.96 | **99.97** | 99.97 | 99.90 | 99.96 | 99.93 | **99.98** | 99.57 | 99.98 | 99.77 | 99.94 | 99.92 | **99.98** | 99.95 | 97.05 |
| INS | **99.01** | 99.92 | **99.46** | 99.85 | 98.94 | 99.37 | 99.15 | 95.23 | 98.04 | **99.93** | 98.98 | **99.86** | 98.55 | 99.81 | 99.17 | 85.05 |
| DUP | **99.90** | **100** | **99.95** | **95.70** | 87.80 | **100** | 93.50 | 84.62 | 75.00 | 100.00 | 85.71 | 84.40 | 97.80 | **100** | 98.89 | 89.78 |
| INV | **99.10** | **100** | **99.55** | 98.28 | 78.80 | 99.49 | 87.95 | 90.61 | 97.20 | 99.59 | 98.38 | **100** | 98.80 | **100** | 99.40 | 85.63 |
| TRA | 98.90 | **100** | 99.45 | **99.11** | **100** | 99.86 | **99.93** | 86.10 | **100** | 98.25 | 99.12 | 96.67 | 99.59 | 99.86 | 99.73 | 64.93 |
| Total | **99.49** | **99.95** | **99.72** | **99.64** | 97.95 | 99.67 | 98.81 | 96.83 | 97.65 | 99.94 | 98.78 | 99.36 | 99.15 | 99.90 | 99.53 | 90.79 |
| ONT |  |  |  |  |  |  |  |  |  |  |  |  |  |  |  |  |
| DEL | **99.98** | 99.91 | **99.95** | **99.96** | 99.90 | 99.58 | 99.74 | **99.96** | 95.02 | **99.93** | 97.41 | 99.95 | 99.94 | 99.92 | 99.93 | 97.06 |
| INS | **99.07** | **99.89** | **99.48** | **99.77** | 98.90 | 99.09 | 98.99 | 95.23 | 93.36 | 99.81 | 96.48 | 99.82 | 98.58 | 99.53 | **99.05** | 85.38 |
| DUP | **99.70** | 99.60 | **99.65** | **96.79** | 87.60 | 99.77 | 93.29 | 86.76 | 69.40 | 99.71 | 81.84 | 89.48 | 95.30 | **99.79** | 97.49 | 89.82 |
| INV | **97.90** | 99.59 | **98.74** | 98.47 | 79.50 | 99.62 | 88.43 | 88.68 | 90.90 | 99.78 | 95.13 | **100** | 97.40 | **99.80** | **98.58** | 89.12 |
| TRA | 98.90 | **100** | 99.45 | **98.90** | **100** | 99.86 | **99.93** | 86.93 | **100** | 98.25 | 99.12 | 97.37 | 99.59 | 99.86 | 99.73 | 66.21 |
| Total | **99.46** | **99.87** | **99.67** | **99.66** | 97.96 | 99.36 | 98.66 | 96.84 | 92.91 | 99.86 | 96.26 | 99.54 | 99.00 | 99.73 | 99.36 | 91.10 |
| PacBio CLR |  |  |  |  |  |  |  |  |  |  |  |  |  |  |  |  |
| DEL | **99.99** | 99.92 | **99.96** | **99.97** | 99.63 | **99.95** | 99.79 | 99.94 | 99.25 | 99.94 | 99.59 | 99.91 | 99.78 | **99.95** | 99.86 | 98.09 |
| INS | **98.48** | 99.34 | **98.91** | **99.76** | 98.05 | 98.98 | 98.51 | 95.51 | 96.71 | 99.83 | 98.25 | 99.47 | 97.71 | **99.89** | 98.79 | 88.03 |
| DUP | **99.60** | 99.70 | **99.65** | **98.90** | 89.20 | 99.78 | 94.19 | 85.87 | 72.10 | 99.86 | 83.74 | 83.50 | 95.70 | 99.90 | 97.75 | 93.21 |
| INV | **99.60** | 99.40 | **99.50** | 99.20 | 85.50 | 99.88 | 92.13 | 84.44 | 97.10 | **100** | 98.53 | **100** | 98.90 | 99.60 | **99.25** | 89.08 |
| TRA | 98.63 | **100** | 99.31 | **98.97** | **100** | 96.43 | 98.18 | 93.83 | **100** | 98.12 | 99.05 | 97.57 | **100** | 99.86 | **99.93** | 65.85 |
| Total | **99.27** | 99.63 | **99.45** | **99.79** | 97.80 | 99.50 | 98.64 | 96.72 | 96.76 | 99.89 | 98.30 | 99.16 | 98.61 | **99.90** | 99.25 | 92.93 |

Rec, recall; Pre, precision; F1, F1 score; GT, genotype concordance. The highest recall, precision, F1 score, and genotype concordance are marked in bold.

In addition to SV detection accuracy, PSVGT consistently delivered superior SV genotyping performance (**Table 1**). PSVGT achieved the highest genotype concordance for deletions, insertions, and duplications on CLR and ONT datasets, and for duplications and translocations on HiFi datasets. Specifically, PSVGT reached genotype accuracies of 98.90% (CLR), 96.79% (ONT), and 95.70% (HiFi) for duplications, outperforming Sniffles2 (84.62-86.76%), cuteSV (83.50-89.48%), and DeBreak (89.78-93.21%). For translocations, PSVGT achieved genotyping accuracies ranging from 98.90% to 99.11% across sequence types, compared with 96.67-97.57% for cuteSV, 86.10-93.83% for Sniffles2, and 64.93-66.21% for DeBreak. While both PSVGT and cuteSV performed well in inversion genotyping, PSVGT achieved higher recall rates and a more balanced trade-off between recall and precision. Collectively, these results demonstrate that PSVGT provide robust and accurate SV detection and genotyping across diverse long-read sequencing platforms, combining high recall, precision, and genotype concordance.

### PSVGT performance on real long-read datasets

We further benchmarked PSVGT against DeBreak, cuteSV, and Sniffles2 using real human long-read data from the HG002 reference sample, comprising 60× PacBio CLR, 50× PacBio HiFi, and 47× ONT datasets. Since high-confidence benchmark call sets for inversions and translocations in HG002 are not available, our benchmarking focused on deletions and insertions. For deletion detection, PSVGT achieved the highest F1 scores across all long-read platforms (95.10-97.52%), outperforming DeBreak (89.95-97.39%), cuteSV (95.42-97.21%), and Sniffles2 (92.94-96.58%), while maintaining high genotype concordance (96.97-98.53%). Insertion calling was generally more challenging across all tools, with lower performance than for deletions. Nevertheless, PSVGT maintained strong performance, achieving recall of 92.65-97.32%, precision of 91.99-94.90%, and F1 scores of 92.32-95.51%, comparable to–and in some cases exceeding–those of DeBreak, cuteSV, and Sniffles2 (**Table 2**). Unlike most SV callers that apply a fixed supporting-read threshold, PSVGT integrates locus-specific supporting-read counts with local read-depth estimates to derive adaptive support criteria. Consistent with simulation benchmarks, PSVGT and cuteSV ranked among the top performers in genotype concordance, with PSVGT achieving 93.68-98.70% compared with 94.61-98.56% for cuteSV (**Table 2**). Taken together, these results closely mirror the simulation-based evaluations and demonstrate that PSVGT achieves a robust balance between recall, precision, and genotyping accuracy, consistently outperforming or matching state-of-the-art SV callers.

**Table 2.**
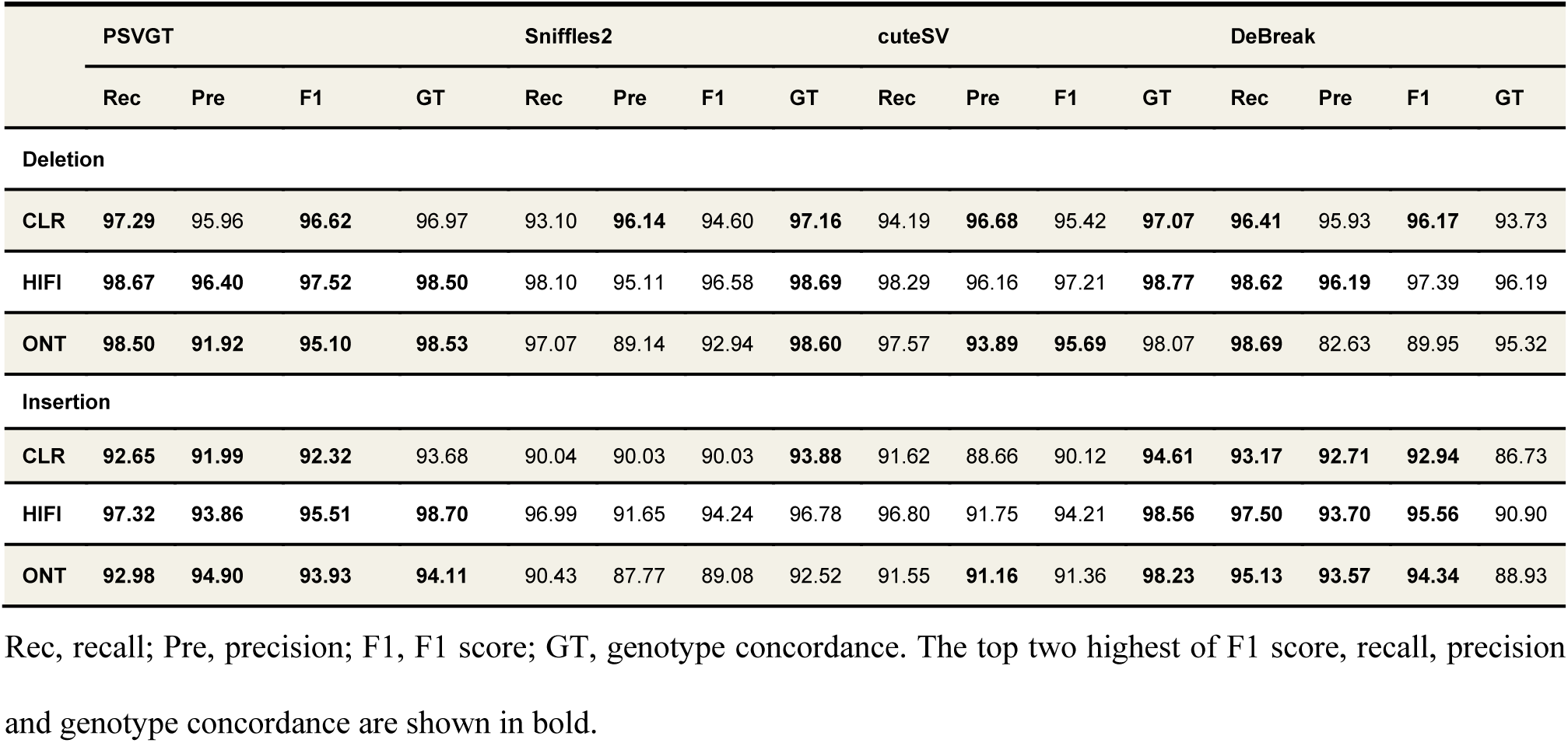
PSVGT performance on the real data of HG002 long-reads sequencing datasets.

### PSVGT performance on downsampled long-read datasets

To assess the effect of sequencing depth on SV detection accuracy and sensitivity, we generated downsampled datasets at 5×, 10×, 15×, 20×, 30×, and 40× coverage using simulated PacBio CLR and ONT reads, as well as PacBio HiFi reads from HG002. As expected, SV detection accuracy improved with increasing sequencing depth across all tools **(Fig. 2)**. Notably, PSVGT achieved ∼90% recall and ∼95% F1 score starting at 10× coverage, demonstrating higher sensitivity than DeBreak, cuteSV, and Sniffles2. At 30× coverage, most tested tools reached saturation in recall and F1 score. Nevertheless, PSVGT consistently achieved the highest performance, with recall and F1 score ranging from 95.75% to 99.39%, followed by DeBreak (95.90-99.20%), Sniffles2 (95.12-98.47%), and cuteSV (95.13-98.19%) **(Fig. 2)**. Importantly, the genotype concordance of PSVGT was largely insensitive to low sequencing depth, maintaining approximately 95% even at 10× coverage. Although DeBreak showed slightly higher recall and F₁ scores than cuteSV and Sniffles2, it exhibited substantially lower genotyping accuracy, particularly at low coverages (<30×), remaining only slightly above 90% even at 50× coverage. Overall, PSVGT consistently outperformed existing tools in both SV detection sensitivity and genotyping concordance, particularly at low sequencing depths, owing to the local depth-adaptive filtering implemented in the KLOOK clustering module, which effectively accommodates uneven read coverage.

**Fig. 2.**
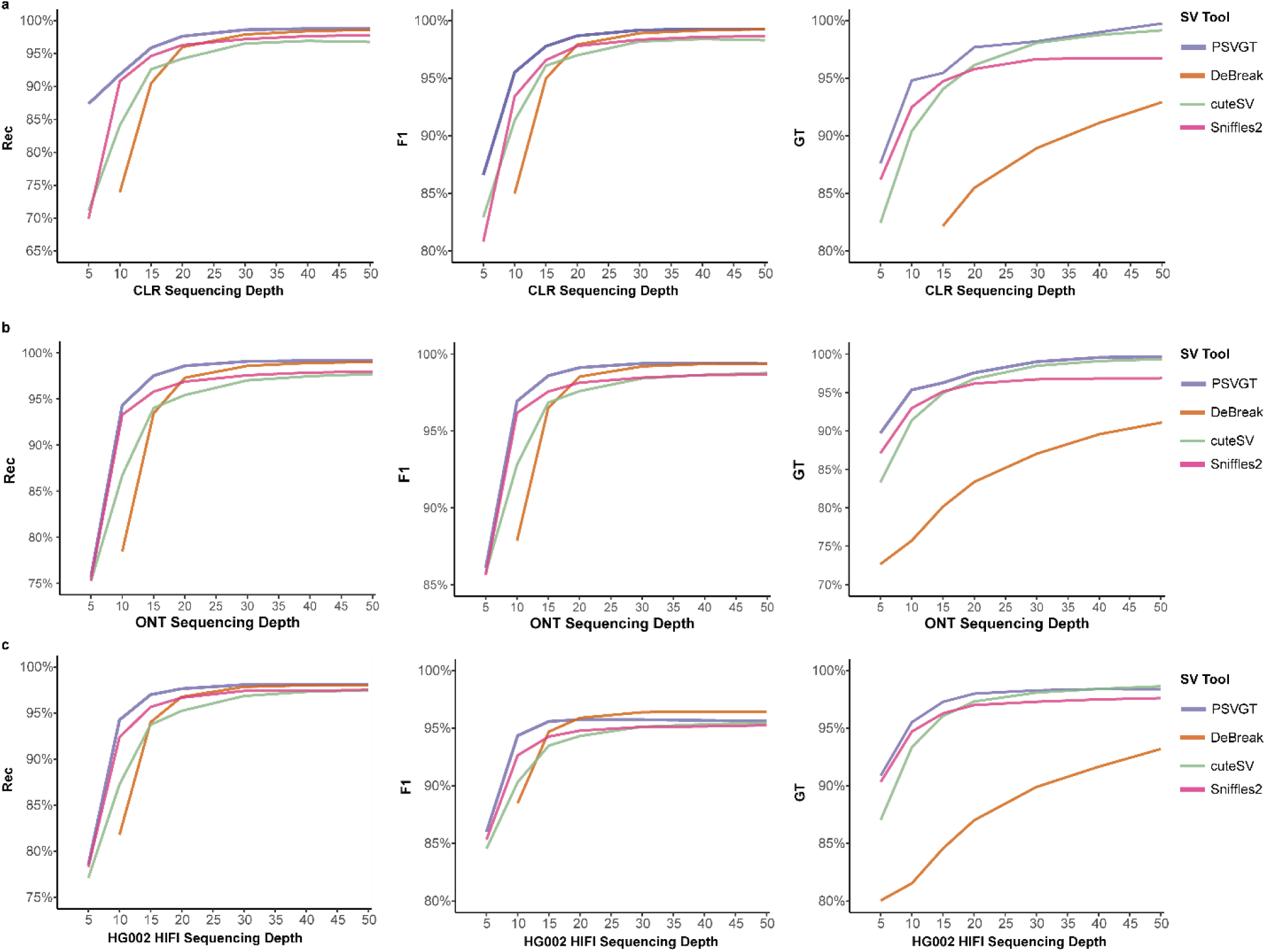
PSVGT performance on downsampled long-read datasets. **a-c**, Comparison of four SV detection tools on downsampled long-read datasets, including simulated PacBio CLR **(a)**, ONT **(b)**, and PacBio HiFi **(c)** datasets. Performance metrics–recall (Rec), F1 score (F1), and genotype concordance (GT)–are plotted against sequencing depth (x-axis).

### Accuracy of SV detection using long-read datasets

SV detection sensitivity and breakpoint accuracy are often constrained by read support and signal density, prompting us to evaluate the performance across size-stratified SVs using long-read datasets. Across the full SV size spectrum, DeBreak and PSVGT consistently outperformed cuteSV and Sniffles2 in terms of F1 score (**Fig. 3a & Fig. S2a & Fig. S3a**). In particular, DeBreak excelled at moderate-length insertions (100-300 bp), whereas PSVGT showed superior performance for large insertions (>2 kb) (**Fig. 3a & Fig. S3a**). This advantage of PSVGT likely arises from the local depth-adaptative filtering implemented in the KLOOK clustering module, which improves SV calling in complex, low-coverage regions and enhances sensitivity to large SVs. For instance, PSVGT successfully detected a 5,071-bp insertion (X:2238538-2238539) supported by only a single segment alignment, which was missed by other state-of-the-art tools (**Fig. S4**).

**Fig. 3.**
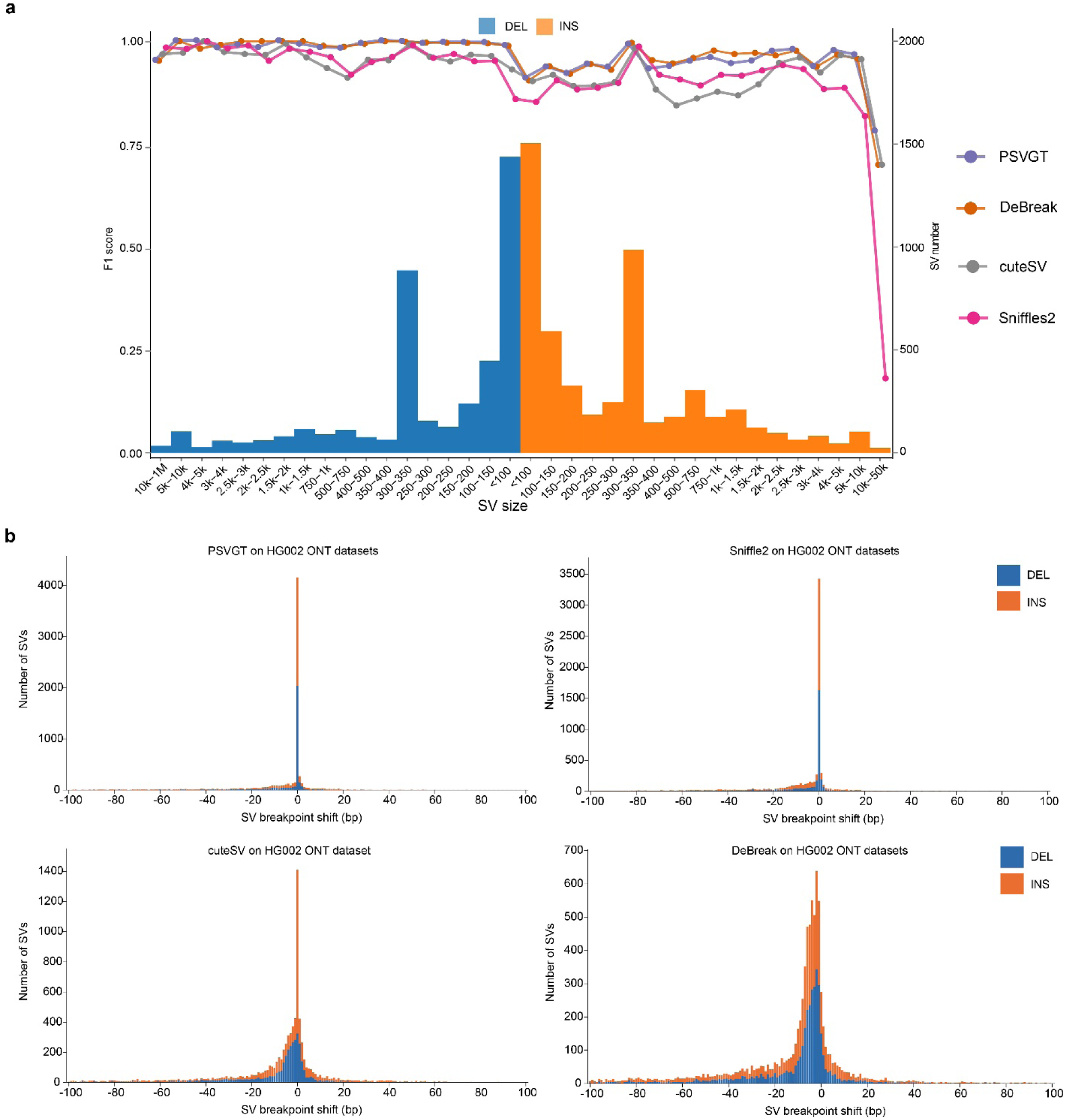
SV detection accuracy using ONT datasets. **(a)**, F1 score distributions across SV sizes for deletions (DEL) and insertions (INS). Histograms indicate the number of SVs within each size bin, whereas lines show F1 scores. **(b)**, Breakpoint shift distributions (≤100 bp) of SVs detected from ONT datasets.

Accurate breakpoint determination is critical for reliable SV genotyping. Using ONT data, PSVGT identified 45.86% (4,159/9,069) of SVs with exact breakpoints and 58.69% (5,323/9,069) within 5-bp shift, substantially outperforming Sniffles2 (38.05%), cuteSV (15.52%), and DeBreak (2.95%) (**Fig. 3b**). This better performance of PSVGT was consistent across HiFi and CLR datasets, where PSVGT exhibited markedly reduced breakpoint shifts compared to other tools (**Fig. S2b & Fig. S3b**). Benchmarking on simulated data further corroborated these findings, demonstrating that the dynamic clustering strategy implemented in PSVGT enables both accurate and efficient breakpoint resolution (**Fig. S5-S7**). Together these results indicate that the dynamic KLOOK clustering algorithm in PSVGT provides highly consistent breakpoint determination and effectively mitigates ambiguity in SV breakpoint detection across diverse long-read sequencing platforms.

### PSVGT performance on haplotype-resolved assembly

The depth-adaptive filtering implemented in PSVGT enables precise resolving of multiple sequence alignments, providing an effective strategy for discovering SVs between reference genomes and haplotype-resolved assemblies. To evaluate PSVGT performance on haplotype-resolved assemblies, we benchmarked it against two state-of-the-art tools, dipcall and SVIM-asm, using both simulated data and real phased assembled genomes. All three tools achieved perfect recall, precision, and F1 scores on simulated benchmarks, as well as genotype concordance (exceeding 99%). In contrast, performance diverged substantially on the real phased assembly (**Table S3**). The dipcall exhibited the greatest decline, with recall and precision dropping to 86.72% and 80.41%, respectively, whereas PSVGT and SVIM-asm maintained high recall rates of 97.48% and 96.43%, respectively. The precision of both PSVGT and SVIM-asm reached ∼92%. Notably, PSVGT achieved the highest genotype accuracy (98.77%), outperforming SVIM-asm (95.54%) and dipcall (92.31%) (**Fig. 4a**). Size-stratified analysis revealed comparable performance among the three tools for deletions, while dipcall underperformed for insertions, with F1 scores falling below 75%, particularly for insertions larger than 10 kb (**Fig. 4b**). This deficiency arises from an algorithmic limitation in dipcall, where insertion lengths are incorrectly summed when the two haplotypes at the same locus harbor insertions of different sizes (**Fig. S8**). In contrast, PSVGT showed a slight advantage in detecting large insertions (>10 kb), achieving an F1 score of 75% compared with 70.59% for SVIM-asm. PSVGT also demonstrated breakpoint accuracy comparable to assembly-specific callers, confirming its sensitivity across both long-read and chromosome-level sequences (**Fig. 4c**).

**Fig. 4.**
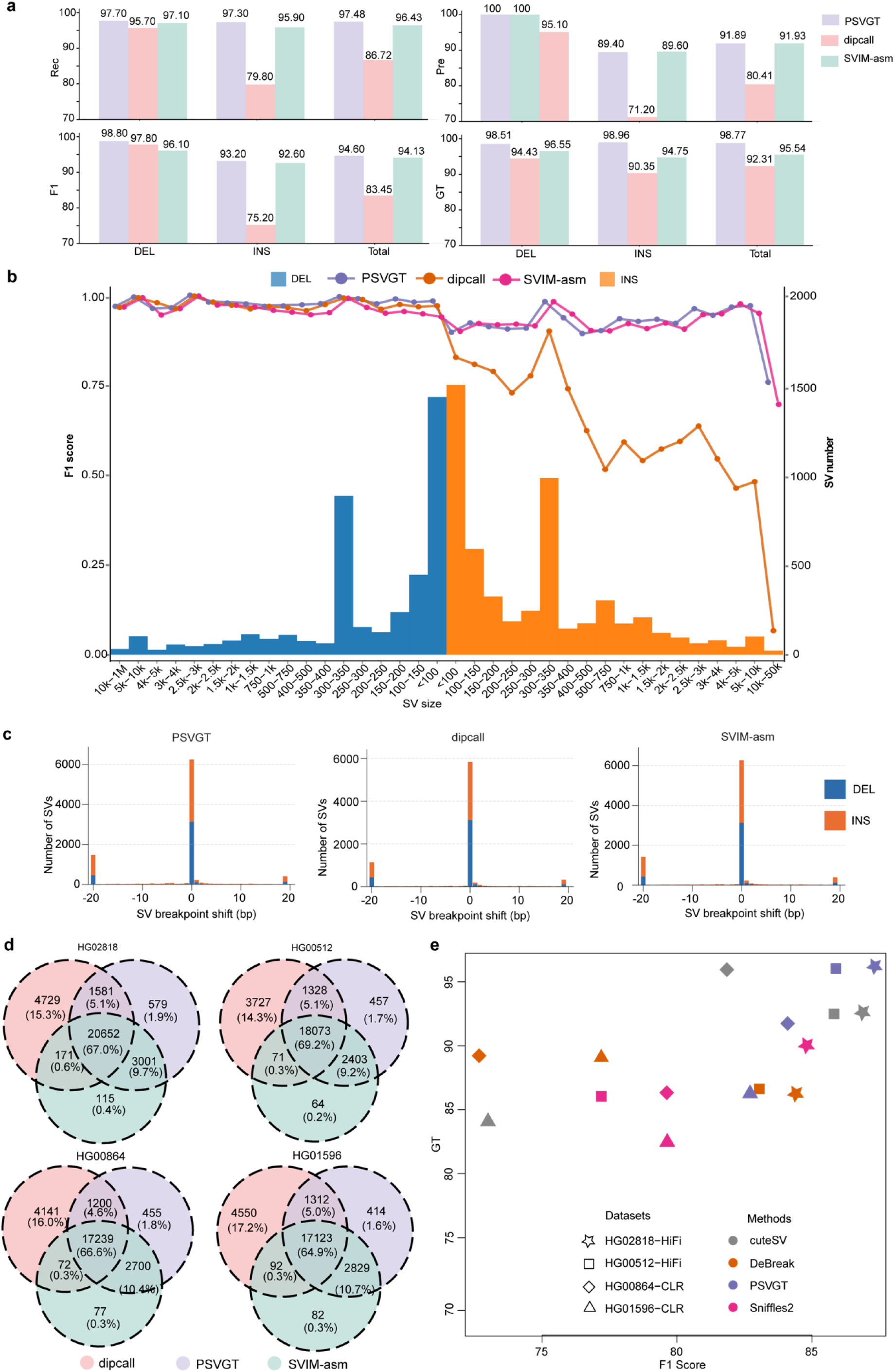
Performance of PSVGT on phased diploid assemblies and long-read datasets. **(a)** Performance of PSVGT, dipcall, and SVIM-asm on HG002 phased assemblies. **(b)** F1 score distributions across SV sizes for phased HG002 assemblies. Histograms showthe number of deletions (DEL) and insertions (INS) within each size bin, while lines indicate F1 scores. **(c)** Breakpoint shift distributions for SVs detected by PSVGT, dipcall, and SVIM-asm. **(d)** Venn diagrams of SV calls among assembly-based methods on phased diploid assemblies, HG02818, HG00512, HG00864, and HG01596. **(e)** Performance of SV discovery tools on HGSVC2 long-read datasets. Scatter plots show F1 score versus genotype concordance (GT) for cuteSV, DeBreak, PSVGT, and Sniffles2 across four datasets.

Since PSVGT is designed to address SV detection and genotyping in both assembled genomes and long-read datasets, we further validated its performance using haplotype-resolved genomes–assembled from PacBio CLR/HiFi reads–from the HGSVC2 project^27^. Intersecting the SV call sets generated by dipcall, PSVGT, and SVIM-asm revealed that 64.9-69.2% of total SVs were constistently identified across samples. PSVGT showed high concordance (75.6-78.4%) with SVIM-asm across all individuals, whereas dipcall produced a substantial number of likely false positives due to the aforementioned insertion-length summation artifact (**Fig. 4d**, **Fig. S8**). Using the intersection of PSVGT and SVIM-asm calls from phased assemblies as a provisional ground truth, we next benchmarked PSVGT against long-read SV callers cuteSV, DeBreak, and Sniffles2 on the corresponding PacBio CLR and HiFi datasets. Across all long-read benchmarks, PSVGT consistently achieved superior performance in both F1 scores and genotype concordance. Notably, PSVGT achieved F1 scores above 85% and maintained genotype concordance exceeding 96% on HiFi datasets (**Fig. 4e & Table S4**). Together, these results demonstrate that PSVGT provides robust and highly accurate SV detection and genotyping across both haplotype-resolved assemblies and long-read sequencing data.

### PSVGT performance on real autotetraploid genome datasets

Given PSVGT’s capacity in supporting diverse sequence types and variable ploidy levels, we further benchmarked its performance using real datasets derived from tetraploid potato. We assessed the performance of PSVGT using fully haplotype-resolved genome assemblies as well as coordinated HiFi datasets at ∼30× coverage per haplotype that were available for three distinct cultivars–Eig, Flo, and BdF. These datasets provided a rigorous benchmark for SV detection because of the complex genomic architecture inherent in tetraploid organisms, in which multiple homologous chromosomes can harbor distinct allelic variants. In this benchmark, PSVGT dynamically adapts the number of clusters according to ploidy, enabling the identification of SV signals from four haplotype assemblies and precise phasing of events such as deletions and insertions that occur on different combinations of homologous chromosomes. For instance, triploid SVs were exclusively detected by PSVGT from HiFi read mapping, as the algorithm effectively utilized local depth adaptation to resolve SV allele dosage (**Fig. 5a**). In haplotype-resolved assemblies, SV genotypes were defined in a phased format such as “1|0|1|0” when the same SV was detected on haplotypes 1 and 3, providing fine-grained resolution of allele dosage and inheritance patterns in polyploid genomes. Across the three potato cultivars, the numbers of SVs identified at different ploidy levels revealed substantial allelic complexity. Diploid SVs, present in two of the four haplotypes, ranged from 8,806 to 18,722 per cultivar. Triploid SVs, in which three haplotypes carried SVs of distinct size, ranged from 447 to 1,761, whereas fully tetraploid SVs–distinct SVs present on all four haplotypes–were comparatively rare, with 12 to 72 events per cultivar (**Fig. 5b, Fig. S9**). To further assess PSVGT sensitivity for polyploid SV detection using long-read data, we treated these diploid, triploid, and tetraploid SVs identified from haplotype-resolved genome assemblies as the ground truth set of multi-allelic SVs (mSVs) (**Table S5, S6, S7**). Although performance was lower than that observed in diploid genomes, PSVGT consistently achieved the highest recall across all three cultivars, outperforming existing long-read SV callers such as cuteSV, Sniffles2, and DeBreak (**Fig. 5b**). The better performance of PSVGT stems from its KLOOK clustering design, which dynamically adjusts cluster numbers for polyploid genomes and utilizes allele-specific local depth-adaptive filtering to recover SVs in complex, low-coverage regions. In contrast, existing SV callers such as cuteSV, Sniffles2, and DeBreak are primarily optimized for autodiploid (human) genomes and exhibit limited sensitivity when applied to polyploid data.

**Fig. 5.**
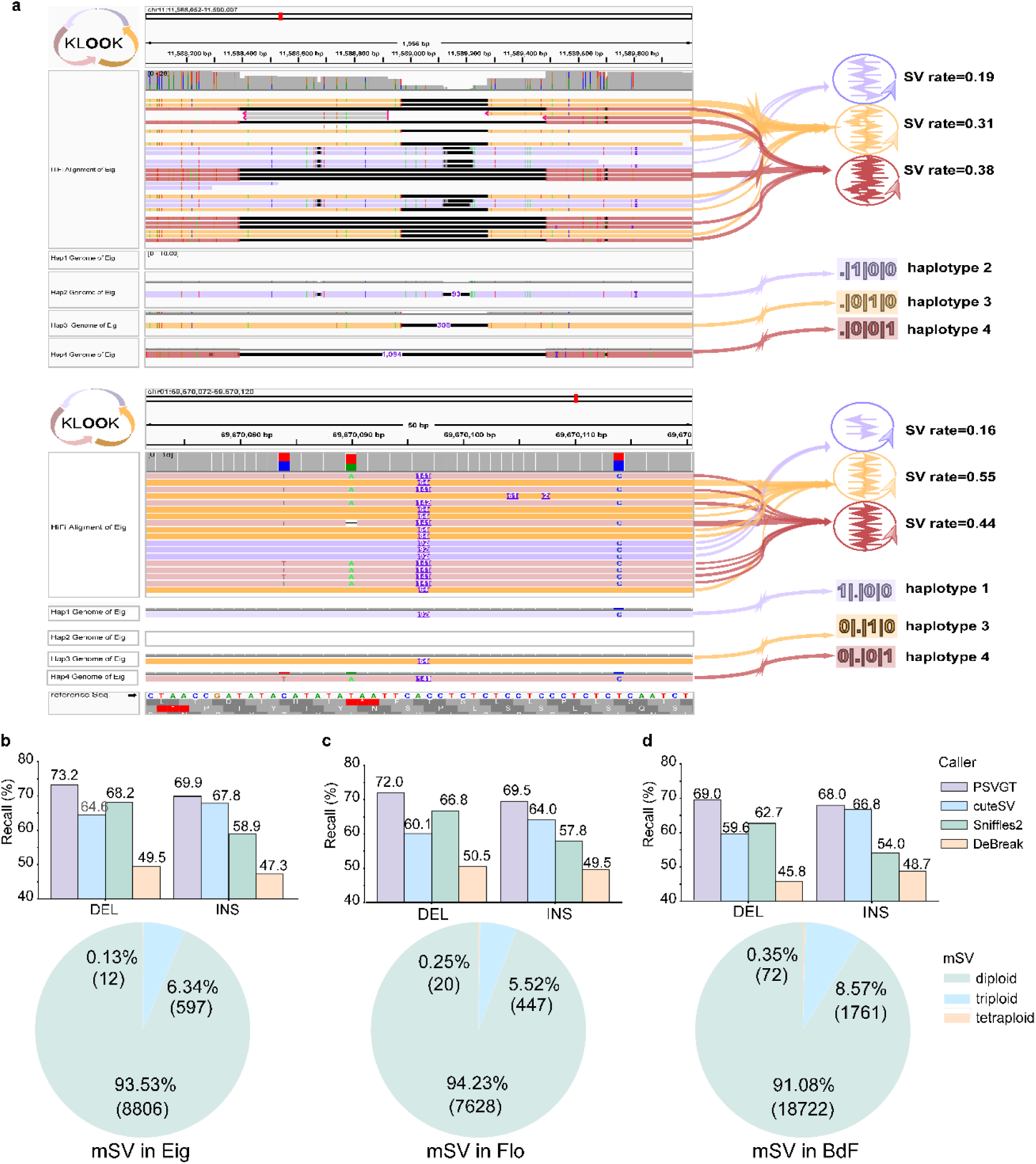
Performance of PSVGT on autotetraploid species. **(a)** Example of a triploid deletion and insertion event in the four-haplotype genome (top) and the corresponding HiFi read alignment (bottom). **(b-d)** Recall performance and distribution of multi-allelic SVs (mSVs) in the tetraploid potato cultivars Eig (b), Flo (c), and BdF (d). Bar plots show mSV recall rates for SV callers, while pie charts indicate the proportions of diploid, triploid, and tetraploid SVs in each cultivar.

### PSVGT performance on real datasets of allopolyploid species

Current long-read SV detection tools are predominantly designed and optimized for diploid genomes, such as humans or inbred model species. However, many important crop plants possess allopolyploid genome architecture. Allopolyploid species, arising from the hybridization of two or more divergent progenitors, present substantial challenges for accurate SV discovery and genotyping. Specifically, high sequence similarity between homologous chromosomes, uneven allele dosage, and frequent inter-subgenome rearrangements can confound alignment- or split-read-based SV detection. To assess PSVGT performance in such complex genomes, we evaluated PSVGT using haplotype-resolved genome assemblies of sour orange (*Citrus × aurantium* (Lour.) Engl.), an allopolyploid species composed of two divergent parental genomes, designated here as hap-M (maternal) and hap-P (paternal). Using the haplotype-resolved assemblies, PSVGT identified 13,968 deletions and 13,422 insertions in the hap-M genome, and 17,327 deletions and 16,897 insertions in the hap-P genome. These SVs corresponded to 22,977 diploid SV loci and 45,954 alternative alleles across the two subgenomes. To evaluate SV detection performance on long-read data, we treated the assembly-derived SVs as the ground truth mSV set and assessed recall across four SV callers. Sniffles2 recovered 13,599 alternative alleles, cuteSV recovered 14,265, and DeBreak recovered 12,357. In contrast, PSVGT recalled 16,254 SV alleles, outperforming the three other tools (**Table S8**). This result demonstrates that PSVGT substantially enhance sensitivity for detecting allelic variation in allopolyploid genomes. In addition to superior recall, PSVGT achieved the highest F1 score and genotyping concordance among the evaluated tools, indicating a favorable balance between precision and sensitivity (**Table S9**). Collectively, these results show that PSVGT not only captures more mSV events in allopolyploid genomes but also provides accurate genotyping of those variants. Taken together, our analyses demonstrate that PSVGT is well-suited for SV discovery and genotyping in allopolyploid plant species, underscoring its potential for population genomics and evolutionary studies in complex non-model plant systems.

### PSVGT performance on population-scale short-read datasets

Population-scale resequencing predominantly relies on short-read datasets; however, accurate detection of insertions from short reads remains a major challenge. We benchmarked PSVGT against two state-of-the-art short-read SV callers, Manta and DELLY, using both simulated and real HG002 short-read datasets. While all three callers showed comparable performance for deletion detection, PSVGT demonstrated clear superiority in insertion detection. In simulated datasets, PSVGT achieved an insertion recall of 94.59%, substantially higher than Manta (67.65%) and DELLY (26.34%) (**Table S10**). In real HG002 short-read data, insertion detection was challenging for all tools due to short read length, heterozygosity and repeats. Nevertheless, PSVGT maintained an insertion recall of 16.65%, approximately twice that of Manta and nearly 20-fold higher than that of DELLY (**Table S10**). To further evaluate PSVGT across different genomic architectures, we assessed its performance in plant species spanning a wide range of repeat content, including *Arabidopsis thaliana* (27% repetitive)^29^, rice (51%)^30^, lettuce (74.2%)^31^, and maize (84%)^32^. We first applied PSVGT to short-read datasets from eight *A. thaliana* ecotypes, with SVs identified by both PSVGT and SVIM-asm from genome assemblies and genotyping results derived from corresponding long-read alignments defined as ground-truth benchmarks (**Table S11**). Across these ecotypes, PSVGT achieved the highest overall recall for indels, with an average recall of 64.62% (58.57-67.86%), substantially outperforming Manta (36.75%; 27.22-43.27%) and DELLY (27.28%; 21.25-32.80%) (**Fig. 6a & Table S11**). Notably, the three tools showed pronounced performance differences in deletion and insertion detection due to their distinct algorithm designs. F1 scores for insertions were consistently lower than those for deletions; however, PSVGT showed only a modest decline compared with DELLY and Manta. Although DELLY achieved the highest precision (91.64%) and genotype concordance (99.26%), its recall for insertions was extremely low (5.66%), resulting in a substantially reduced F1 score of approximately 10.62%. In contrast, PSVGT achieved a balanced performance, with 56.59% insertion recall, 88.88% precision, and 94.91% genotype concordance (**Fig. 6a & Table S11**). Importantly, PSVGT’s performance on short reads (recall >62%, precision >87%, F1 score >73%, and genotype concordance >94%) closely mirrored that of long-read data. This comparable performance likely reflects PSVGT’s strategy of assembling short reads into draft contigs with lengths comparable to long reads for SV detection, thereby effectively recapitulating long-read signals for SV detection and genotyping. In addition, we assessed breakpoint accuracy by by refining genotype (PSVGT_srGT) using short reads pre-aligned BAM and found that SV breakpoints supported by short-read alignments were highly consistent with those inferred by PSVGT (**Fig. 6a)**.

**Fig. 6.**
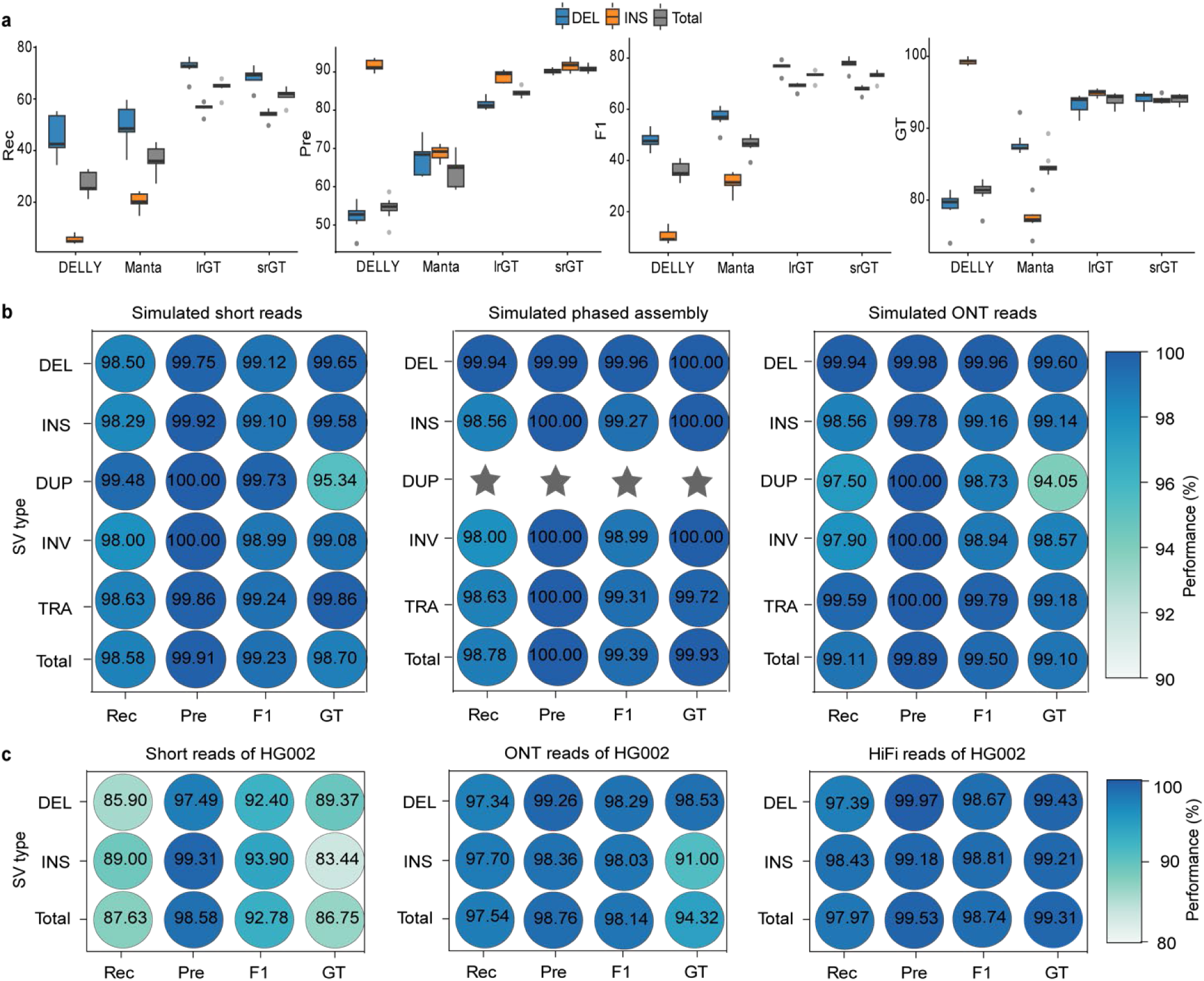
Performance of PSVGT on short-read datasets and SV genotyping. **(a)** Comparison of SV detection metrics across different tools. Boxplots show F1 score (F1), genotype concordance (GT), precision (Pre), and recall (Rec) for DELLY, Manta, lrGT (genotyping based on Short-read–derived contigs alignment) and srGT (genotyping based on short-read alignment), stratified by deletions (DEL), insertions (INS), and all SVs combined (Total). **(b, c)** Genotyping performance on simulated data **(b)** and real data of HG002 **(c)**. Performance metrics include recall (Rec), precision (Pre), F1 score (F1), and genotype concordance (GT) across different SV types and sequencing platforms.

Similar trends were observed in rice, maize, and lettuce short-read datasets, in which PSVGT consistently outperformed Manta and DELLY in both recall and precision, particularly for insertion detection **(Fig. S10, Table S12-S14)**. Collectively, these results demonstrate that the *de novo* assembly-based strategy implemented in PSVGT substantially improves insertion detection from short-read data, outperforming current state-of-the-art tools and exhibiting robust performance across plant genomes with different repeat content. This contrasts with the local reassembly strategy employed by Manta and the split-read/paired-end mapping approaches used by DELLY, highlighting PSVGT’s versatility for population-scale SV discovery.

### Force-genotyping of PSVGT

Most existing SV genotyping tools rely on individual evidence signals to assign genotypes. In contrast, PSVGT is designed to enable population-scale SV detection and genotyping, with genotyping modules specifically engineered to support force-genotyping of predefined SVs across large populations without the computational burden of whole-genome SV discovery. To evaluate force-genotyping module of PSVGT, we applied it to 21,843 candidate SVs derived from a simulated PacBio HiFi dataset and augmented this set with 11,700 synthetic false positives generated by shifting SV coordinates by 50 kb. These SVs were genotyped using 50× short reads, simulated diploid assemblies, and ONT reads. By leveraging candidate signals from the HiFi dataset, PSVGT achieved superior genotyping performance across short-read, phased assembly, and ONT datasets, maintaining recall above 97% and genotype concordance approaching or exceeding 99% across all SV types (**Fig. 6b**). For short-read datasets, genotype concordance reached 99% for most SV types, with the exception of duplications, which showed slightly reduced concordance (94.05-95.34%) due to inherent alignment limitations.

We further benchmarked SV genotyping on the real HG002 dataset using 8,906 high-confidence SVs and 6,416 synthetic false positives, using 30× short-read, 47× ONT-read, and 50× HiFi-read datasets. Deletions were genotyped more accurately than insertions, with long-read datasets achieving recall above 97% and genotype concordance exceeding 98% for most SV types. In contrast, on 30× short-read data PSVGT achieved an overall recall rate of 87.63% and genotype concordance of 86.75% (**Fig. 6c**). The reduced performance on HG002 relative to simulated datasets is primarily due to genuine genomic divergence from the reference genome, including abundant SNPs and small indels that can potentially compromise alignment accuracy.

Integrating long- and short-read sequencing is commonly used for population-scale SV discovery and genotyping, offering a balance between accuracy and cost. Using 17,895 non-redundant SVs identified from eight *Arabidopsis* ecotypes as the ground truth, we applied the force-genotyping module of PSVGT to population-scale short-read datasets. Overall, genotyping performance exceeded that observed in the HG002 benchmarks. Notably, insertion genotype concordance (88.69-90.83%) was comparable to or higher than deletion concordance (84.85-87.25%), a pattern that contrasts with the HG002 results (**Table S15**). This difference likely reflects the largely homozygous nature of *Arabidopsis* ecotypes, whereas human samples exhibit substantially higher heterozygosity. Collectively, these results demonstrate that PSVGT enables rapid and robust SV genotyping across sequencing platforms, featuring high recall and accurate genotyping in population-scale short-read datasets.

### SV annotation and marker development module in PSVGT

SVs can influence genome function through impacts on gene bodies, transcript architecture, and gene expression. To facilitate functional interpretation of detected SVs, PSVGT incorporates an annotation module that classifies putative functional impacts of SVs based on their genomic context. SVs are categorized into the following functional classes: SVExp (overlaps with UTRs or promoters potentially affecting gene expression), GenePAV (presence/absence variation caused by loss of coding sequences), IntronExten/Shorten/Loss (intronic structural alterations), Frameshift, and GeneAS (disruption of canonical splice sites leading to alternative splicing). Applying this annotation framework to *Arabidopsis thaliana* ecotypes (Db-1 versus Col-0) revealed that 80.9% of detected SVs were intergenic and the remaining 19.1% were predicted to have potential functional impacts, with 2.5% classified as SVExp, 3.8% as Frameshift, 5.8% as intronic changes (IntronExten/IntronShorten/IntronLoss), 5.4% as GeneAS, and 1.6% as GenePAV (**Fig. 7a**). These annotations provide valuable insight into the functional relevance of SVs and offer a basis for downstream prioritization in studies of gene regulation, trait association, and genome evolution.

**Fig. 7.**
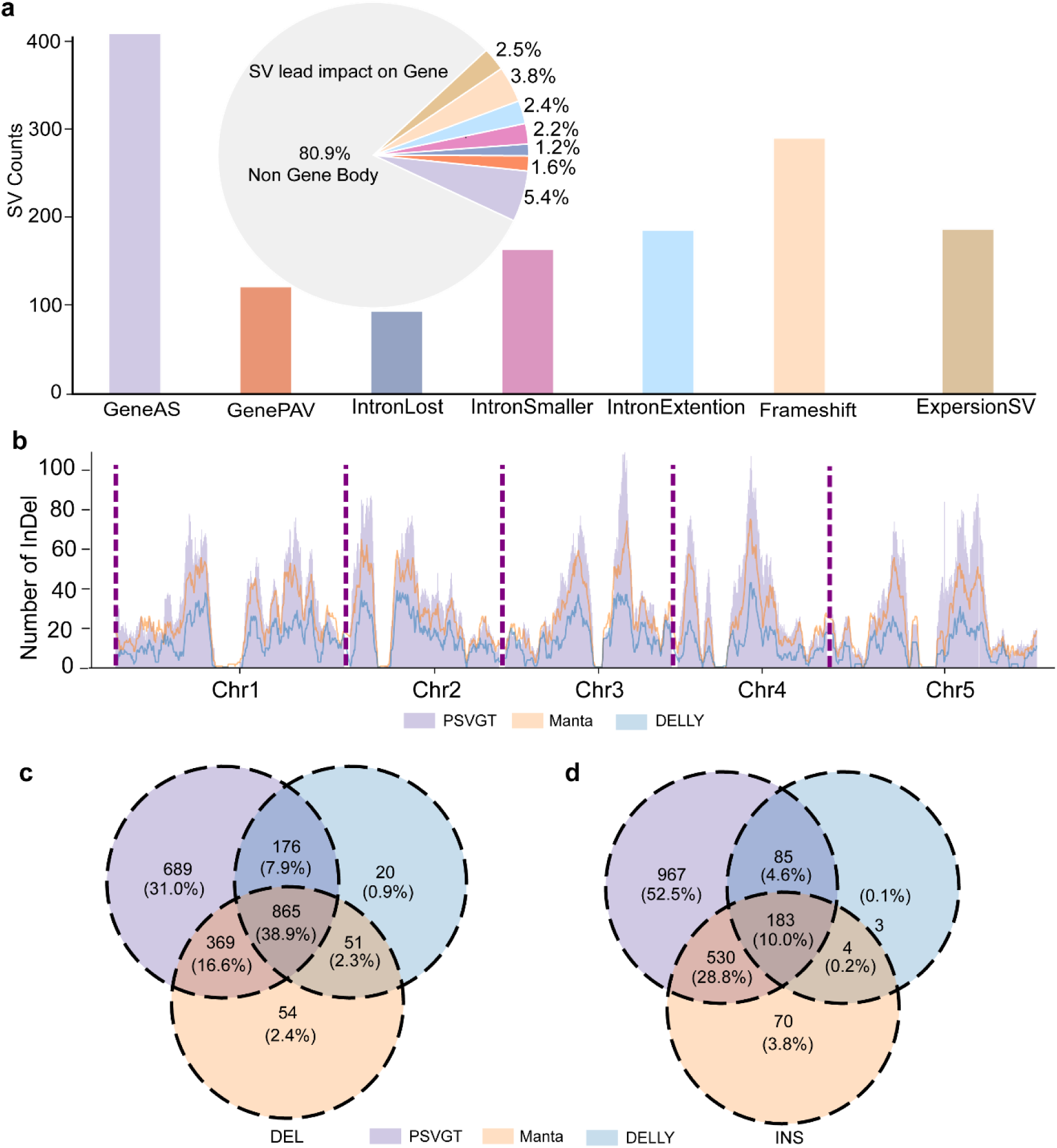
SV annotation and marker development module of PSVGT. **(a)** Example annotation of SVs in the PSVGT module. Annotation features are represented by distinct colors. The bar plot indicates the number of SVs annotated under each representative term, while the pie plot displays the proportion of each annotation category. **(b)** Density distribution of 50–600 bp InDels across *Arabidopsis* chromosomes. Density curves, color-coded for tools (PSVGT, purple; Manta, orange; DELLY, blue) show the density of InDel number across chromosome1–5. (**c-d)** Intersection of InDel calls among tools. Venn diagrams showing the number of deletions **(c)** and insertions **(d)** identified by PSVGT, Manta, and DELLY across all eight datasets.

Molecular markers are widely used in population genetics and breeding, particularly for gene mapping and cloning. To further facilitate the application of SVs in genomic and genetic analyses, PSVGT integrates a marker-development module-- SVInDelPrimer, enabling indel marker development by selecting user-defined SV size ranges and designing primers from flanking sequences. Using eight *Arabidopsis* short-read datasets, PSVGT consistently outperformed Manta and DELLY in detecting InDels of experimentally optimal sizes (50-600 bp), enabling genome-wide, high-density marker development (**Fig. 7b**). For deletion markers, PSVGT uniquely identified 689 candidates from a shared core set of 865, compared with fewer than 55 unique deletions detected by either Manta or DELLY (**Fig. 7c**). For insertion markers, PSVGT reported 1,765 candidates, including 967 unique events (**Fig. 7d**), whereas Manta identified 787 candidates (90% overlapping with PSVGT) and DELLY identified 275 candidates (97% overlapping with PSVGT). The high intersection rate of SV markers indicates that PSVGT captures a broader and reliable set of SV markers from short-read datasets. Primer design for candidate SV markers in PSVGT is performed using the primer3-py API, yielding optimized, high-confidence PCR primers for downstream experimental validation.

### Runtime and memory usage

To assess runtime and memory efficiency, we compared PSVGT with other long-read SV callers using the 50× pre-aligned PacBio HiFi BAM file from HG002. All analyses were conducted on a high-performance server equipped with an Intel Xeon Gold 6230R CPU (104 cores at 2.10 GHz) and 512 GB RAM, using 10 threads. PSVGT completed SV detection—including split-read and depth signal extraction, local depth-adaptive filtering, ploidy-aware KLOOK clustering, and genotyping—in 35 min with a peak memory usage of 3.9 GB. In comparison, cuteSV completed in 25 min using 5.1 GB RAM, DeBreak in 20 min using 10.2 GB RAM, and Sniffles2 in 15 min using 2.3 GB (**Table S16**). The modest increase in runtime for PSVGT reflects its more comprehensive filtering and clustering strategies, which improve sensitivity in regions with uneven or complex coverage.

We then evaluate runtime and memory usage using 50× human short-read dataset. *De novo* assembly using MEGAHIT required 2.5 h and a peak memory of 139.4 GB. Following assembly, PSVGT detected and genotyped SVs from the assembled contigs in 8.5 min using 3.0 GB RAM. In comparison, Manta processed the pre-aligned BAM file in 14 min with peak memory of 0.9 GB, whereas DELLY required 97 min (only supporting single thread) and 6.0 GB RAM (**Table S17**). The genotyping phase of PSVGT, which assigns allelic states to 25,561 SVs based on short reads pre-aligned BAM file, completed in under 3 min with a peak memory of 0.9 GB (**Table S16**). Overall, PSVGT completed SV discovery and genotyping in ∼35 min on a standard 50× long-read human dataset and in ∼8 min on assembled contigs. These results indicate that advanced filtering and clustering features based on ploidy in PSVGT do not introduce excessive computational overhead, making it well suited for high-throughput SV analysis in diploid and polyploid genomes, as well as population-scale studies.

## Discussion

Structural variation is a major contributor to genomic diversity and disease etiology, exerting complex functional effects by directly altering gene structures and cis-regulatory elements and indirectly perturbing long-range regulatory interactions. Despite its importance, accurate SV discovery, genotyping, and interpretation remain challenging, particularly in species with complex genome architectures. Existing SV callers are typically optimized for specific sequencing technologies or data types: DELLY and Manta are designed for short reads; cuteSV, Sniffles2, and DeBreak focus on long reads; and dipcall and SVIM-asm rely on genome assemblies. In contrast to these specialized SV callers, PSVGT provides a unified framework that integrates short-read, long-read, and whole-genome assemblies datasets, achieving performance that matches or exceeds state-of-the-art tools across all data types. Notably, PSVGT overcomes key limitations of short-read-based SV detection–especially for insertions–by adopting a *de novo* assembly strategy that achieves higher precision and recall than Manta and DELLY. By integrating complementary sequencing modalities, PSVGT leverages the sensitivity of long reads and assemblies while remaining compatible with widely available short-read data, enabling the recovery of SVs that are routinely missed by conventional short-read pipelines.

Polyploid genomes comprise multiple homologous subgenomes that may each harbor distinct SV alleles, and allopolyploids combine divergent parental haplotypes, greatly complicating sequence alignment, SV discovery, and genotype inference. A major innovation of PSVGT is its local depth-adaptive framework, which is specifically designed to handle complex genome regions and polyploid species. In contrast to most existing SV callers, which implicitly assume diploidy and are unable to estimate allele dosage beyond two haplotypes, the KLOOK clustering algorithm in PSVGT dynamically infers multiple SV clusters based on user-specified ploidy. This design enables accurate resolution of multi-allelic SVs. By extending SV genotype inference to arbitrary ploidy levels, PSVGT provides robust SV discovery and genotyping in both autopolyploid and allopolyploid species, which are common among crop plants.

SV analysis at the population scale requires both high sensitivity and computational efficiency. The force-genotyping module implemented in PSVGT enables genotyping of predefined SV sets across thousands of samples without reprocessing whole-genome SV signals, thereby facilitating population-scale SV genotyping from diverse sequencing data types. In the *Arabidopsis* population benchmark, force-genotyping achieved 87.81% genotype concordance with HiFi-derived genotypes, highlighting its suitability for large-scale population studies. Combined with SV functional annotation, PSVGT provides an integrated framework for post-GWAS analyses and functional genomics studies. Moreover, the comprehensive SV sets inferred by PSVGT can be directly incorporated into pangenome graph construction tools such as vg^33^ and Minigraph^34^. Such integration is expected to reduce reference bias and improve genotype accuracy in pangenome analyses, particularly for species with extensive genomic structural diversity.

Despite the robustness of PSVGT across diverse sequencing data types and species, limitations remain when analyzing short-read data in highly repetitive genomic regions. First, as with most SV callers, PSVGT faces challenges in resolving SVs located within highly repetitive sequences, primarily due to that short-read-derived contigs often fail to assemble accurately in these regions. Additionally, incorporation of poorly aligned short reads in such regions can lead to missed SV calls and erroneous SV genotyping. Second, although the KLOOK algorithm restricts clustering to 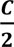 comparisons (where C is the number of aligned SV signal reads) within a target window to efficiently resolve multi-allelic events, regions with extreme read depth–such as duplication-rich or high-copy-number loci–may still experience modest runtime overhead. Nevertheless, KLOOK remains an essential component of PSVGT due to its critical role in accurately resolving alleles, particularly in complex polyploid genomes where multi-allelic SVs are prevalent. Importantly, the integrated processing of hybrid data types in PSVGT–including short reads, long reads, and assemblies–substantially mitigates these challenges by improving assembly continuity and spanning repetitive regions. Together, these capabilities advance SV discovery and genotyping in complex genomes, reinforcing the utility of PSVGT for genomic research, population studies, and crop improvement.

## Materials and Methods

### Raw SV signal detection

To enable SV detection across diverse sequencing data types, PSVGT employs distinct strategies for short- and long-read datasets. For short-read datasets, PSVGT implements a *de novo* assembly-based workflow using MEGAHIT (v1.2.9)^22^ to reconstruct contigs prior to variant calling. For long-read datasets, raw FASTQ files are aligned directly to the reference genome using minimap2 (v2.24)^23^. PSVGT then extracts SV signatures from both intra- and inter-alignments (**Fig. 1a**). Discordant alignment orientations and clipped read positions are used to detect deletions, insertions, duplications, and inversions. Each raw SV signal records the start and end coordinates, SV length, supporting read identifiers, mapping quality scores, and associated sequences. For contig and genome assembly datasets, insertions and deletions are inferred from gaps or extensions within alignments. For long-read datasets, deletions are inferred when reference segments are skipped between adjacent alignment blocks, while insertions are deduced from additional read sequences located between alignment blocks. Duplications are identified by overlapping alignments on the reference genome, and inversions are detected when paired segments align in opposite orientations. Translocations are identified from supplementary alignments (“SA” tags in BAM files) in which a single read maps to distinct chromosomes. All detected SV signals are subsequently clustered and refined using the KLOOK clustering algorithm.

### Dynamic KLOOK clustering and local depth-adaptive filtering

To reduce background noise and handle varying signal densities across the genome, PSVGT implements a dynamic sliding-window clustering strategy coupled with local depth-adaptive filtering. Coordinates of each window are automatically adjusted according to the inferred SV length as defined follows:

SV length ≤ 100 bp: window = (SV_start_ - 200 - 0.2 × SV length, SV_end_ + 200 + 0.2 × SV length);

100 bp < SV length ≤ 500 bp: window = (SV_start_ - 400 - 0.2 × SV length, SV_end_ + 400 + 0.2 × SV length);

500 bp < SV length ≤ 1000 bp: window = (SV_start_ - 600 - 0.2 × SV length, SV_end_ + 600 + 0.2 × SV length);

SV length > 1000 bp: window = (SV_start_ - 1000, SV_end_ + 1000).

Within each genomic window, SV signals are first grouped by variant type and sorted by genomic coordinates. These signals are then clustered using the KLOOK algorithm, where k is set to the sample ploidy (e.g., 2 for diploid, 3 for triploid, and 4 for tetraploid). For each window, the signal will be aggregated through C/2 (C = total signals within the window) iterative comparisons, leveraging SV position, SV length, and read identifier. The top K clusters are retained as putative alleles corresponding to the ploidy level. Each allele from KLOOK is subsequently filtered based on the local coverage of the 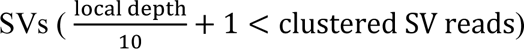.

### Genotyping modules

PSVGT incorporates two specified genotyping modules, each tailored to unique characteristics of data types. The long-read genotyping module, applicable to long-read sequencing data, contigs, and assembled genomes, leverages two complementary sources of evidence for robust genotype inference: (1) local SV signal density and (2) breakpoint coverage information. In contrast, the short-read genotyping module, designed exclusively for short-read datasets, relies exclusively on breakpoint coverage derived from original short-read alignments (rather than assembly) as the primary evidence for genotype assignment.

#### Breakpoint validation

A candidate SV breakpoint is considered valid if the distance between the predicted SV boundary and clipped read ends (left/right) satisfies the following criteria:

For long-read dataset:

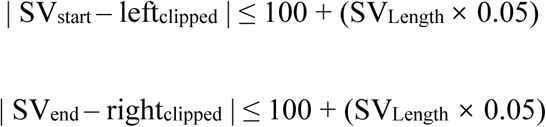

For short-read dataset:

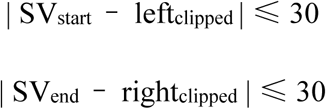

Where SV_start_ and SV_end_ denote the predicted start and end of a candidate SV, left_clipped_ and right_clipped_ represent the clipped end positions of aligned reads, and SV_length_ is the length of candidate SV. This dynamic threshold account for inherent variability in long-read alignment acround breakpoints.

#### Breakpoint covered reads quantification

Reads are classified as breakpoint-covered reads if their alignment spans the breakpoint region with sufficient flanking sequence, as defined as below:

For long-read dataset:

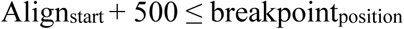

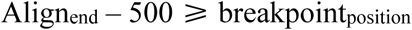

For short-read dataset:

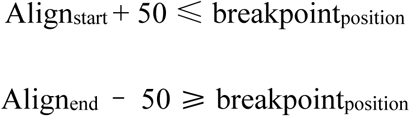

Where Align_start_ and Align_end_ are the start and end coordinates of the read alignment, and breakpoint_position_ refers to either the left or right breakpoint coordinate of the SV. The 500 bp flanking requirement for long reads ensures that reads fully span the breakpoint region with high confidence.

#### Score calculation

Genotypes are inferred based on normalized scores, calculated as the ratio of supporting reads to local genomic depth defined as average read coverage within 1 kb window flanking the SV:

The SV allele score is calculated as:

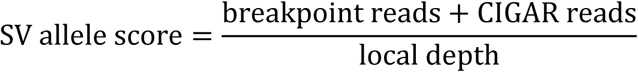

Where breakpoint reads is the number of reads passing breakpoint validation, CIGAR reads is number of reads with CIGAR strings indicating SV-specific alignment patterns. For short-read datasets, CIGAR-based evidence is excluded due to read length limitations.

The reference allele score is calculated as:

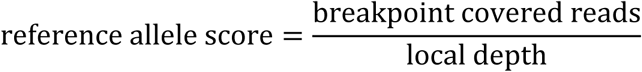

Where breakpoint-covered reads are reads satisfying breakpoint coverage criteria and supporting the reference allele.

### Benchmark metrics

To evaluate the performance of each tool, we calculated recall, precision, F1 score and genotype concordance.

Recall was defined as:

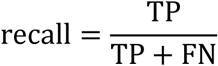

where TP (true positive) refers to SVs detected by the tool that are concordant with the true set, and FN (false negative) refers to SVs present in the true set but not detected.

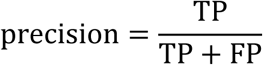

where FP (false positive) refers to SVs detected by tool but not present the in true set.

F1 score was defined as:

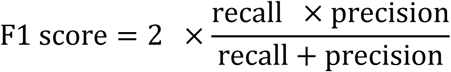

Genotype concordance was defined as:

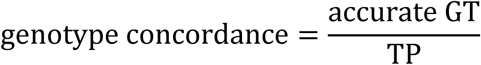

where accurate GT refers to true SVs whose genotypes are concordant with those in the true set.

### Dataset simulation

We simulated SVs consisting of 10,000 deletions, 10,000 insertions, 1,000 duplications, and 1,000 inversions using scripts implemented in DeBreak, with all variants introduced into two independently simulated haploid human genomes. Additionally, 365 translocation sites were collected from published cuteSV datasets^10^. The size distribution of each SV type was consistent with that observed in human genomes. The overall ratio of heterozygous to homozygous events was set to 2:1, with alleles randomly assigned to the hs37d5 reference haplotype genome. For translocations, VISOR (v1.1.2.1)^35^ was used to generated both heterozygous and homozygous translocated genomes. PacBio CLR datasets were simulated using PBSIM3 (v3.0.4)^24^ with the parameters “--errhmm ERRHMM-RSII.model --depth 25”. ONT and PacBio HiFi reads were generated via Badread (v0.4.1)^36^ with parameters “--quantity 25× --junk_reads 0 --random_reads 0 --glitches 0,0,0” and parameters “--quantity 25× --error_model pacbio2021 --qscore_model pacbio2021 --glitches 0,0,0 --junk_reads 0 - -random_reads 0 --chimeras 0”, respectively. For all datasets, the sequencing depth was uniformly set to 25× per haploid genome. Short reads were simulated using InSilicoSeq (v2.0.1)^26^ with options “--model Novaseq”, generating a sufficient number of reads to achieve an average sequencing depth of 50×.

#### Benchmark on long-read datasets and assembled genome

Minimap2 (v2.24) was used to align PacBio CLR, PacBio HiFi, and ONT reads to the reference genome, with parameters selected for each sequencing platform, as detailed in the minimap2 manual. Sniffles2 (v2.6.0)^9^, cuteSV (v2.1.1)^10^, and DeBreak (v1.2)^12^ were used to call SVs, with the minimum number of supporting reads set from 3 to 10, and the highest F1 score of each tool was used for benchmarking. Only SVs > 50 bp were retained for benchmarking. The haploid genomes were aligned to the reference genome using minimap2 with parameters “-a -x asm5 --cs -t8”. dipcall (v0.3)^17^ and SVIM-asm (v1.0.3)^18^ were then applied to detect SVs from the two simulated haploid genomes with default parameters. Final variant files in VCF format generated by PSVGT and other SV callers were evaluated against the ground truth SV sets to compute precision, recall, F1 score, and genotype concordance. For simulated data, a detected SV was considered a true positive if its breakpoint was within 1 kb of the corresponding ground-truth variant and its length was within 80-120% of the true SV size. Genotype concordance was recorded when the SV genotype matched the ground truth set. For real datasets, SV calling performance was evaluated using Truvari (v5.1.1)^28^ with parameters --passonly -r 1000 -p 0.00 --dup-to-ins. High-confidence insertion and deletion variants from a publicly available benchmark dataset were used as the truth set. Insertions, deletions, and duplications detected by each tool were extracted and compared against the truth set using Truvari. The precision, recall, F1 score, and genotype concordance were obtained from the Truvari reports.

For the four samples from Human Genome Structural Variation Consortium 2 (HGSVC2) resource, SV detection was performed on genome assemblies using PSVGT, SVIM-asm, and dipcall, and on the corresponding long-read datasets using PSVGT, cuteSV, DeBreak, and Sniffles2. Intersection SV sets derived from phased assemblies were used as ground truth to assess the performance of long-read SV detection. For each sample, we defined a series of minimum supporting long reads ranging from 4 to 15 for SV detection and selected the optimal results from these tools for comparison with the results of PSVGT. All call sets were benchmarked against the ground truth using Truvari with the flags --passonly -r 1000 -p 0.00 --no-ref, generating precision, recall, F₁ scores and genotype-concordance metrics for each.

To assess the sequencing depth effect on PSVGT, simulated PacBio CLR reads, ONT reads, and HiFi reads were downsampled to depths of 5×, 10×, 15×, 20×, 30×, 40×, and 50× using samtools (v 1.15.1)^37^. Comparative callers (DeBreak, cuteSV, Sniffles2) were executed with a series of minimum support reads based on the depth of datasets, whereas PSVGT was performed using default configuration. Each downsampled call set was benchmarked against the corresponding ground truth SV set using Truvari, generating precision, recall, F₁ score, and genotype concordance metrics.

### Benchmark on tetraploid and allodiploid genome

For tetraploid potato, haplotype-resolved assemblies and corresponding PacBio HiFi reads were aligned to DM_1-3_516_R44 (version 6.1) using minimap2. mSVs were inferred from the genome alignment of the haplotype-resolved genomes when SVs were detected on different haplotypes with a size fold change ≥ 0.8. A custom script was used to search mSVs from HiFi-derived VCF file to identify regions harboring at least two alternative alleles, which were taken as true recall. The number of clusters in PSVGT was set to four to reflect the tetraploid genome architecture of potato.

For allodiploid citrus, diploid genome assemblies were aligned to sweet orange reference genome (SWO, version 3)^38^. The diploid SVs were inferred from the diploid genome alignments. The same genomic region with SV size fold change ≥ 0.8 was defined as mSV. SV calls derived from PacBio HiFi data using Sniffles2, cuteSV, DeBreak, and PSVGT were used to benchmark. The minimum number of supporting reads was varied from 4 to 10, and the highest F1 score of each tool was used for performance assessment.

### Benchmark on short-read datasets

Short-read–derived contigs and original short reads were aligned to reference genome using minimap2(v2.24)^23^ and bwa (v0.7.17)^39^, respectively. To establish the ground-truth variant set for benchmarking short-read performance, were first identified from haplotype-resolved assemblies using SVIM-asm and the long-read module of PSVGT. Genotypes for these SVs were subsequently assigned by intersecting the calls of cuteSV and PSVGT on the corresponding ONT/HiFi datasets. This genotyped SV set was used as the benchmark for assessing the performance of DELLY, Manta and short-read module of PSVGT. A detected SV was considered a true positive if its predicted breakpoint was located within 1 kb of a ground-truth variant and its length was within 80–120% of the true SV size. Genotype concordance was recorded when the genotype call matched the genotype in the ground-truth set.

### Force genotyping

To evaluate the genotyping performance, SV calls derived from HiFi data using PSVGT were used as the candidate set for genotyping. These SVs were subsequently genotyped in simulated short-read datasets using the short-read genotyping module of PSVGT, and in diploid-resolved genome alignments using the long-read genotyping module. For the *Arabidopsis* population, a high-confidence truth set was constructed by jointly calling SVs from HiFi/ONT reads using the long-read module of PSVGT (parameters: “-lr_homo_rate 0.75 -lr_ref_rate 0.15”) and cuteSV. SV calls from the two methods were merged, and redundant variants were removed to generate a nonredundant SV set. This set was then used as the reference for forced genotyping across all short-read samples using the short-read module of PSVGT (parameters: “-homo_rate 0.3 -ref_rate 0.15”). Genotype concordance was assessed by comparing the forced-genotyping outputs against the truth set.

### Marker module of PSVGT

The SVInDel_primer module enables high-throughput identification of experimentally tractable InDels and produce size-distinguishable PCR amplicons for reference and variant alleles. The Primer design is performed using the primer3 package implemented in Python^40, 41^. For each candidate InDel locus, SVInDel_primer extracts reference sequences franking the predicted breakpoint and designs forward and reverse primers from these regions. Primer design was conducted using uniform global constraints, including an optimal primer length of 21 bp (rang 18-25 bp), optimal melting temperature of 60 °C (range 53–65 °C), GC content 40–80%, and stringent limits on self-complementarity and primer–primer interactions. No ambiguous bases were permitted in primer sequences. Primer pairs are subsequently evaluated to identify breakpoint-spanning combinations that minimize amplicon length while maintaining primer quality and specificity, ensuring efficient amplification of InDels compatible with standard PCR assays.

## Data availability

The HiFi passed assemblies, HiFi, ONT, PacBio CLR, and Illumina sequencing datasets are available at GIAB and NCBI. The high confidence benchmarks of HG002 are available at https://ftp-trace.ncbi.nlm.nih.gov/giab/ftp/data/AshkenazimTrio/analysis/NIST_SVs_Integration_v0.6/HG002_SVs_Tier1_v0.6.vcf.gz. The high confidence regions is available at https://ftp-trace.ncbi.nlm.nih.gov/giab/ftp/data/AshkenazimTrio/analysis/NIST_SVs_Integration_v0.6/HG002_SVs_Tier1_v0.6.bed. The phased assemblies and long read datasets of HGSVC2 are at https://ftp.1000genomes.ebi.ac.uk/vol1/ftp/data_collections/HGSVC2/release/v1.0/assemblies/. The raw illumina, HiFi, ONT sequencing of the 8 ecotypes of *Arabidopsis* were downloaded at EMBL-ENA under the accession number PRJEB62038. And the genome assemblies of the 8 ecotypes of *Arabidopsis* were downloaded at NCBI-SRA under the accession number PRJNA1033522. The rice assembled genomes, long- and short-read datasets were downloaded at EMBL-ENA under accession number PRJEB73710, and T2T rice genome NIP was downloaded from http://www.ricesuperpir.com/web/download. The lettuce short-read datasets were downloaded at China National Center for Bioinformation under accession number CRA014517. The maize short-read datasets were downloaded at EMBL-ENA under accession number PRJEB31061. The haplotype assemblies of the potato genome were downloaded from https://doi.org/10.5281/zenodo.14053896, and the raw sequencing data were downloaded from NCBI-SRA under the accession number PRJNA1074690. The diploid genome assemblies of sour orange were downloaded from http://citrus.hzau.edu.cn/download.php, and raw sequencing datasets were downloaded at NCBI-SRA under the accession number PRJNA1003804. All the datasets used in our study are listed in **Table S1**.

## Code availability

All code is available at https://github.com/lgbTime/PSVGT under the MIT License.

The key custom scripts used in the manuscript are available at https://github.com/lgbTime/PSVGT_Benchmarks_Code.

## Acknowledgements

This research was supported by the National Key Research and Development Program of China (2023YFF1000100), the scientific research start-up funding from Hubei Hongshan Laboratory award no. 11020102, the “Young Scientist Fostering Funds for the National Key Laboratory for Germplasm Innovation & Utilization of Horticultural Crops”. The computations in this paper were run on the bioinformatics computing platform of the National Key Laboratory of Crop Genetic Improvement, Huazhong Agricultural University.

## Contributions

X.W. and Z.F. conceived and led this work. G.L and X.W. designed the framework. G.L. implemented the framework. G.L. and X.L. performed data analysis. G.L., X.W., and Z.F. wrote the manuscript.

