## Supplemental Fig for "A sensitive and accurate framework for population-scale structural variant discovery and genotyping across sequence types"

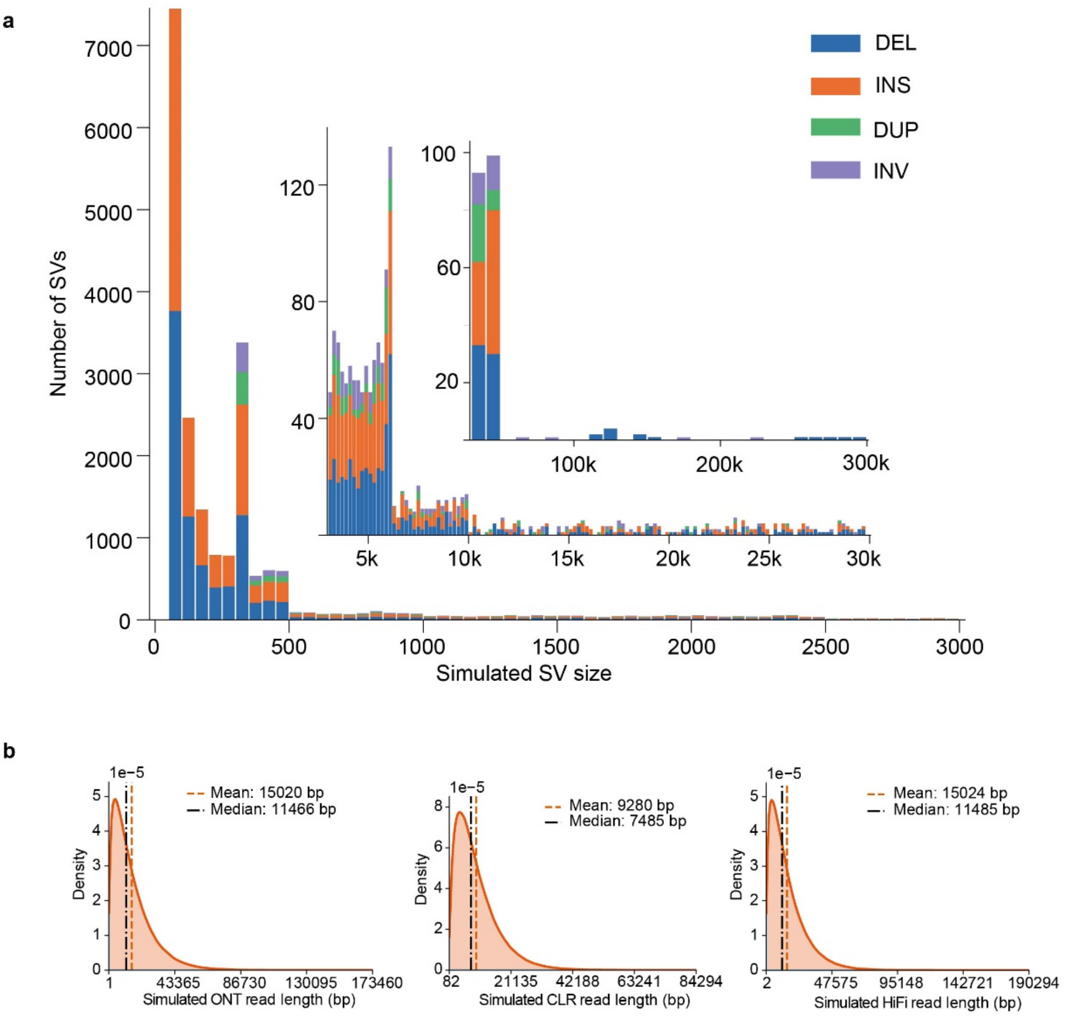


**Figure S1**. **Simulated SVs and read length distribution**. **(a)** The size distribution of simulated SVs. The peak at 300–400 bp range corresponds to simulated *Alu* elements, while the peak at ~6 kb represents mobile elements. **(b)** Read length density distributions of simulated ONT, CLR, and HiFi datasets.

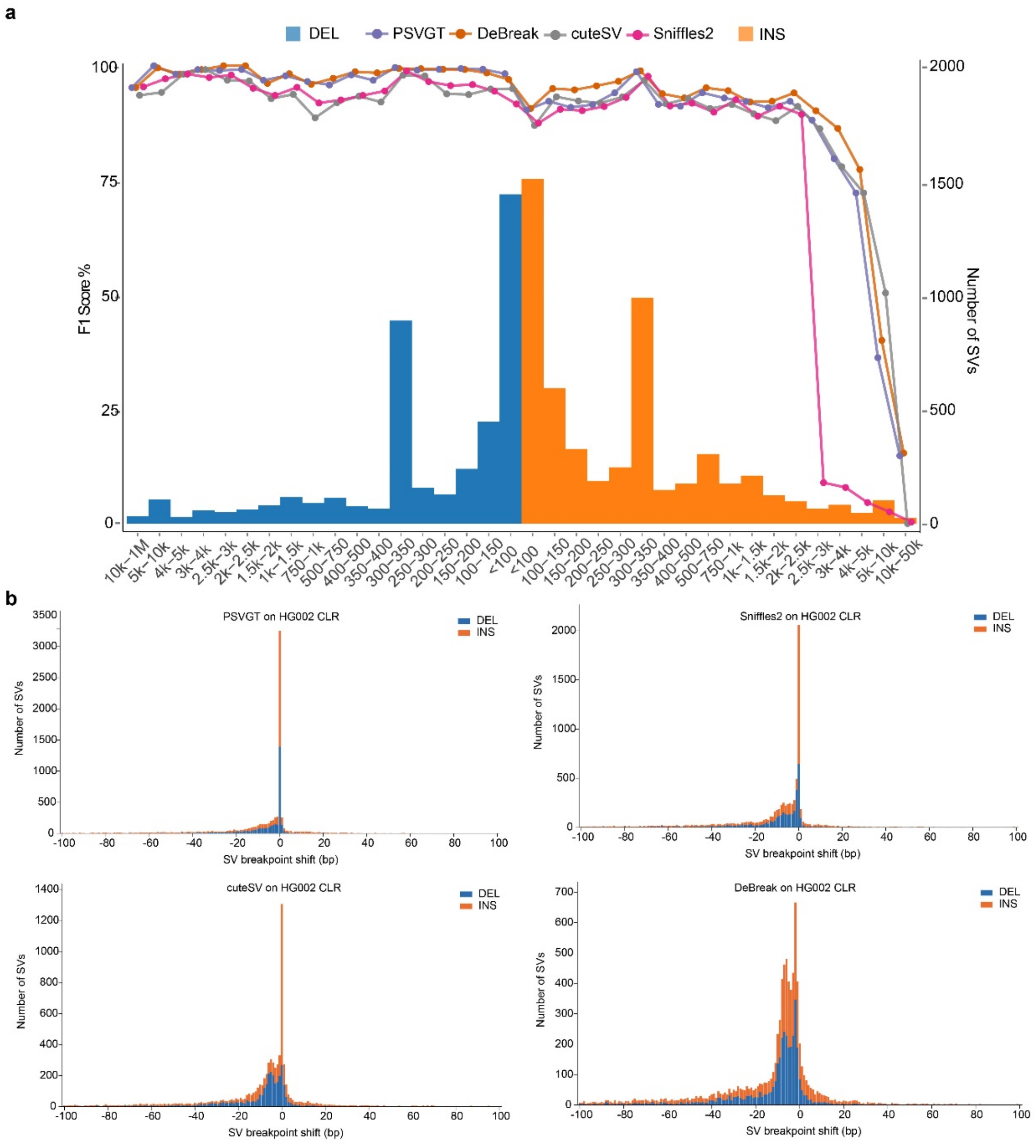


**Figure S2**. **SV detection accuracy on PacBio CLR datasets of HG002**. **(a)** F1 score distribution across different SV size ranges and plotted as colored lines for each evaluated tool. **(b)** SV breakpoint accuracy for the four tools on HG002 CLR datasets, SVs with breakpoint shift ≤ 100 bp were included.

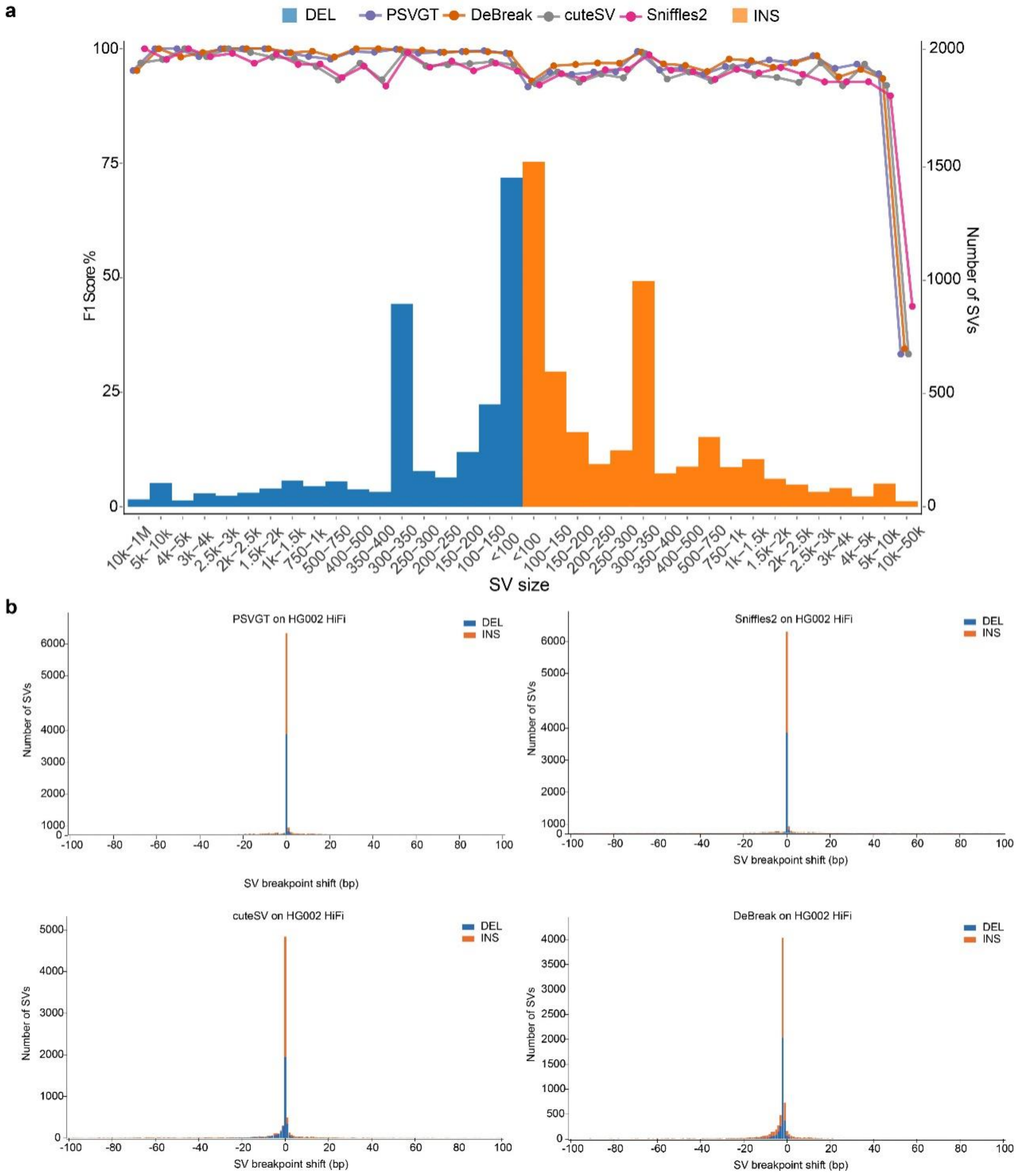


**Figure S3**. **SV detection accuracy on PacBio HiFi datasets of HG002**. **(a)** F1 score distribution across different SV size ranges and plotted as colored lines for each evaluated tool. **(b)** SV breakpoint accuracy for the four tools on HG002 HiFi datasets, SVs with breakpoint shift ≤ 100 bp were included.

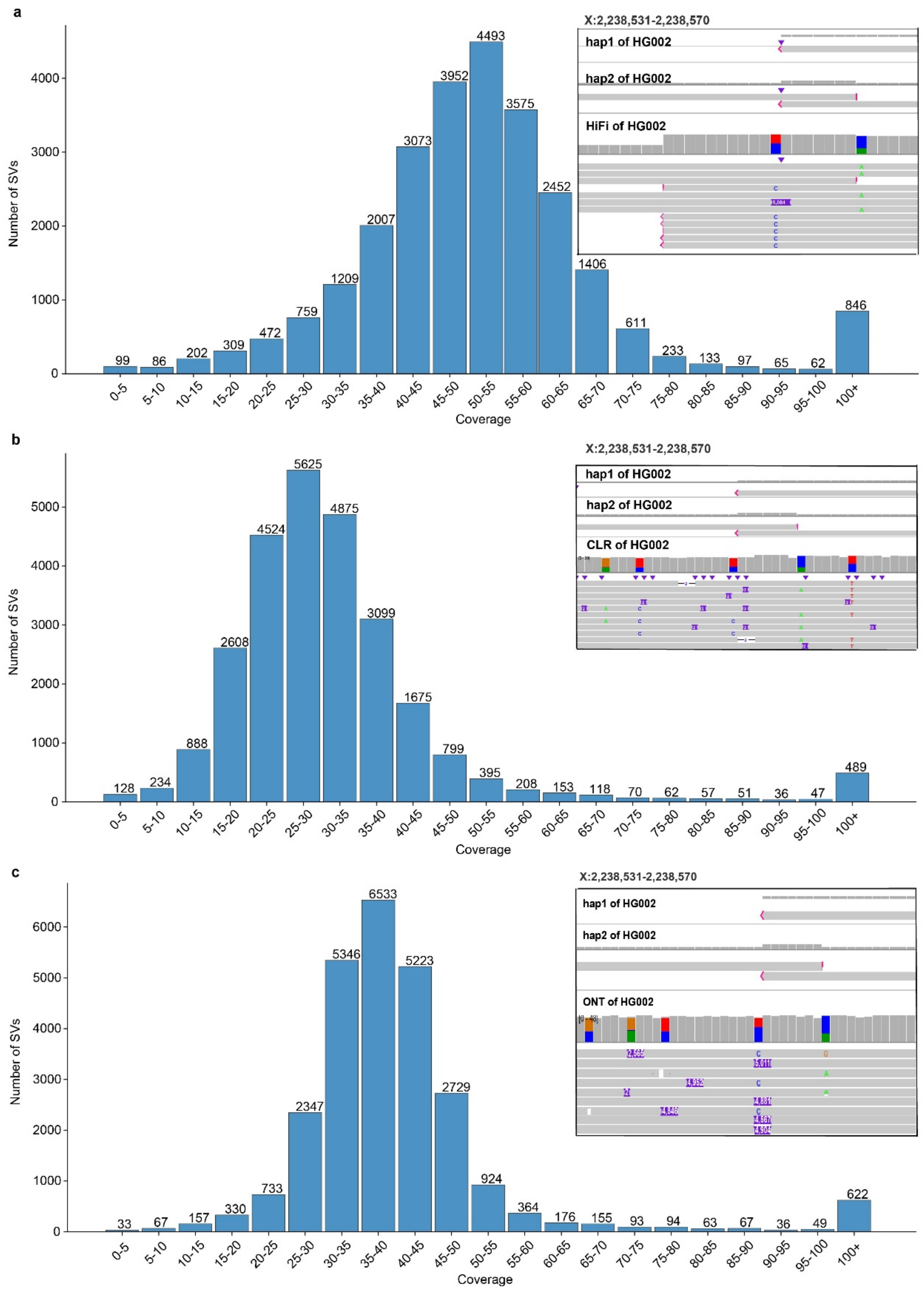


**Figure S4**. **Local coverage distribution of SVs in HG002.** Local coverage profiles of SVs in **(a)** PacBio HiFi, **(b)** PacBio CLR, and **(c)** ONT datasets. IGV snapshots illustrate a 5,071 bp insertion at locus X:2,238,531–2,238,570. Alignment breakpoints are marked in red, and insertion-related CIGAR operations are highlighted in purple. Panels **(b)** and **(c)** show the same genomic region as in **(a)**.

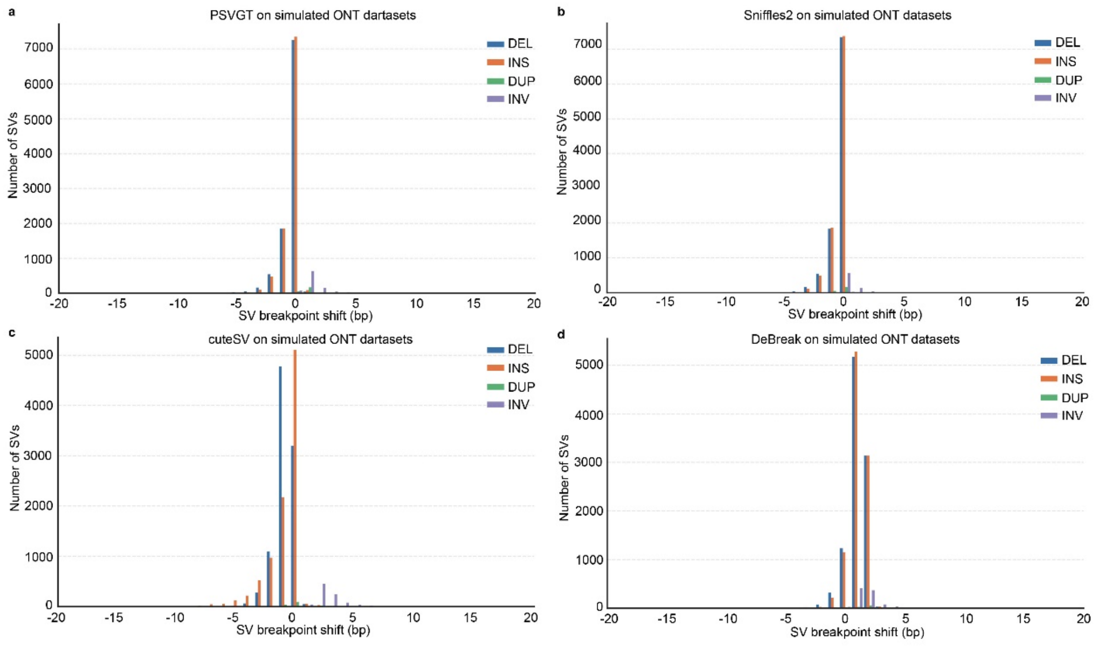


**Figure S5**. **Breakpoint accuracy of SVs on ONT datasets.** SVs with breakpoint shift ≤ 20 bp were displayed. Different SV types are represented by distinct colors. Panels show results for each caller: **(a)** PSVGT; **(b)** Sniffles2; **(c)** cuteSV; **(d)** DeBreak.

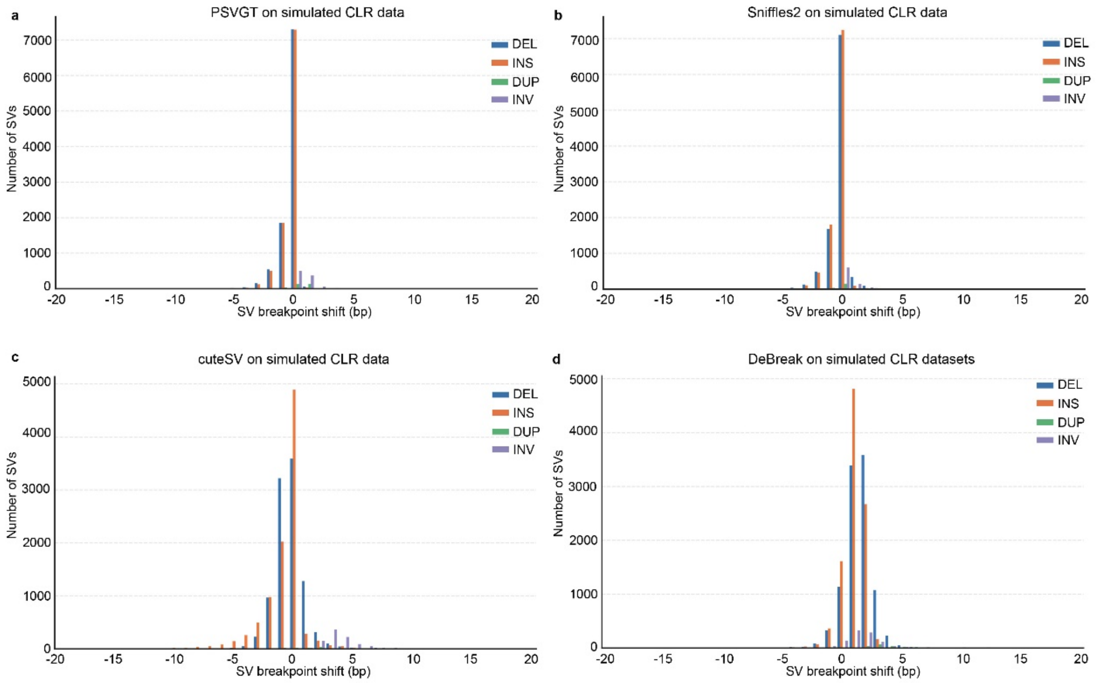


**Figure S6**. **Breakpoint accuracy of SV on CLR datasets.** SVs with breakpoint shift ≤ 20 bp were displayed. Different SV types are represented by distinct colors. Panels show results for each caller: **(a)** PSVGT; **(b)** Sniffles2; **(c)** cuteSV; **(d)** DeBreak.

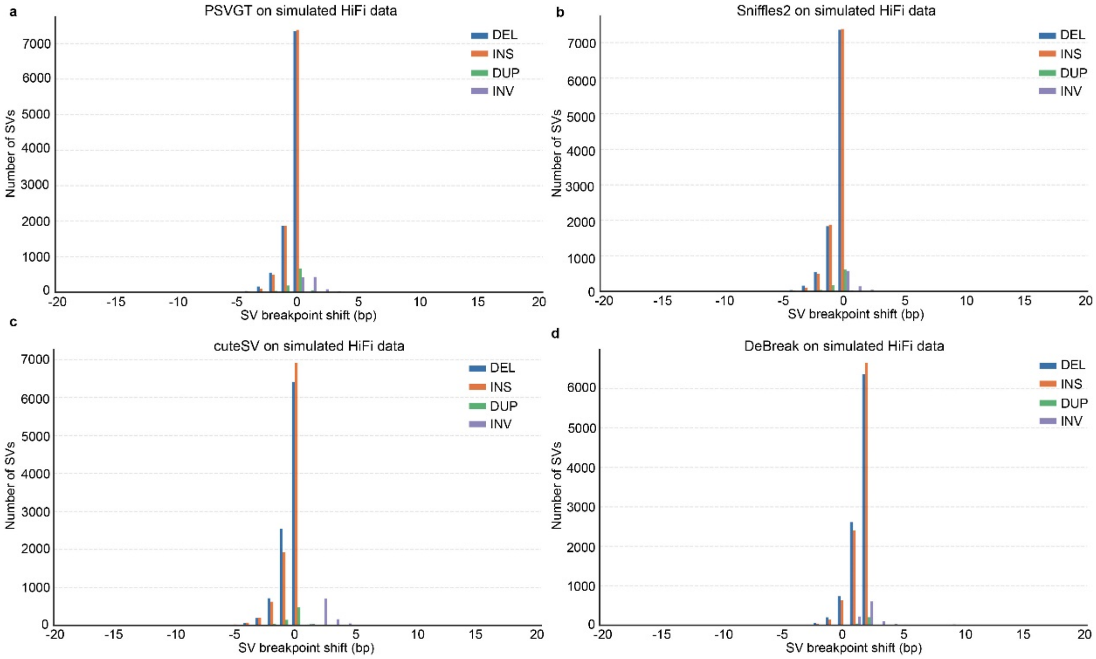


**Figure S7**. **Breakpoint accuracy of SV on HiFi datasets.** SVs with breakpoint shift ≤ 20 bp were displayed. Different SV types are represented by distinct colors. Panels show results for each caller: **(a)** PSVGT; **(b)** Sniffles2; **(c)** cuteSV; **(d)** DeBreak.

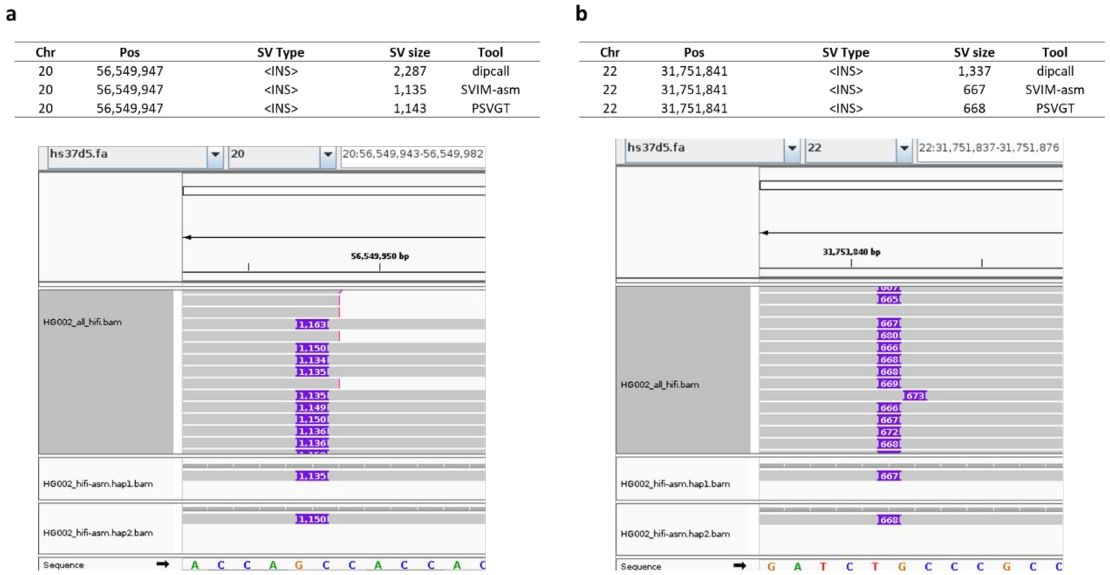


**Figure S8. Comparison of insertion detection on phased diploid assemblies**. **(a)** Insertion calls on chromosome 20 as reported by dipcall, SVIM-asm and PSVGT, visualized in IGV. **(b)** Insertion calls on chromosome 22 as reported by dipcall, SVIM-asm and PSVGT, visualized in IGV.

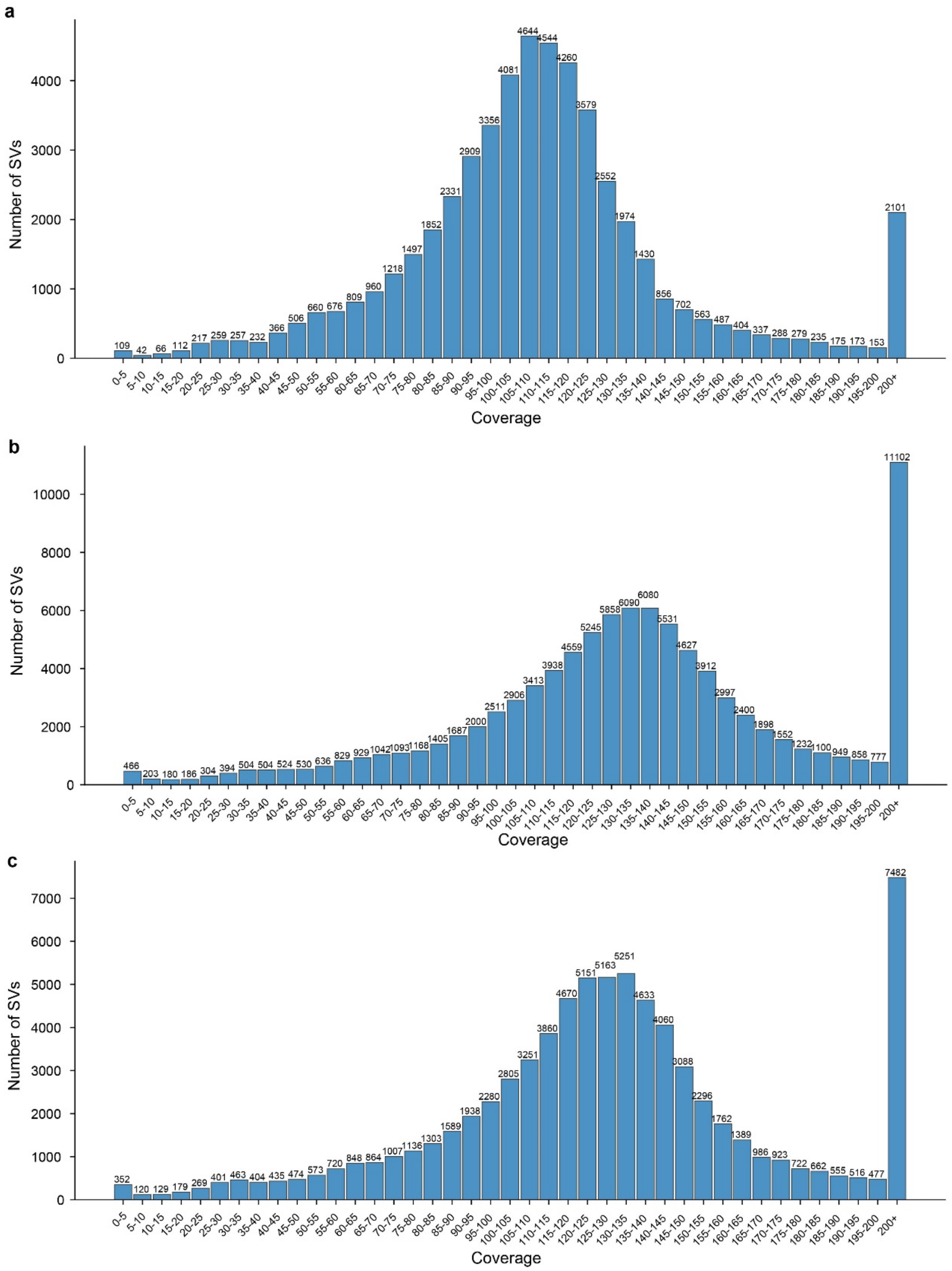


**Figure S9 | Local coverage distribution of SVs in tetraploid potato samples.**
**(a–c)** Histograms showing the local coverage distributions of SVs in PacBio HiFi datasets for Eig **(a)**, Flo **(b)**, and BdF **(c)**. All panels depict the number of SVs across coverage intervals (x-axis: coverage; y-axis: SV count).

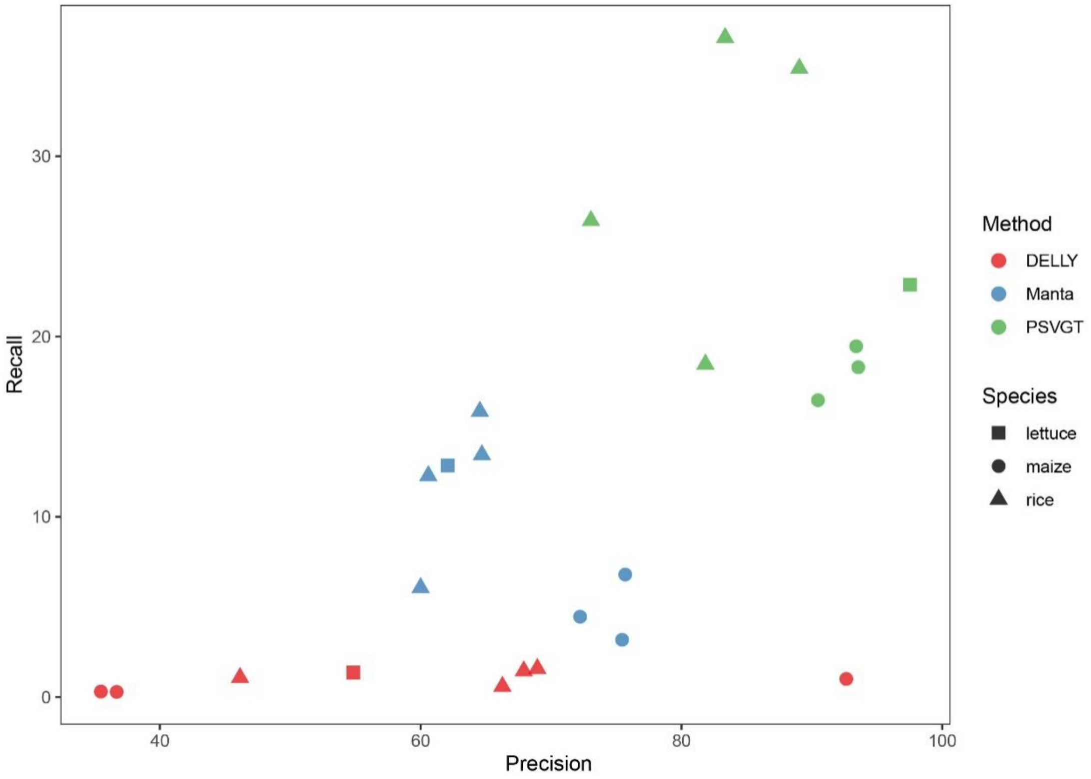


**Figure S10 Comparison of insertion performance across short-read datasets**. A total of eight datasets were included: four from rice, three from maize, and one from lettuce. Different species are represented by distinct shapes, while the performance of each tool is indicated by distinct colors. Insertions from the haplotype genome were used as the ground truth sets for insertions.
